# The impact of learning on perceptual decisions and its implication for speed-accuracy tradeoffs

**DOI:** 10.1101/501858

**Authors:** André G. Mendonça, Jan Drugowitsch, M. Inês Vicente, Eric DeWitt, Alexandre Pouget, Zachary F. Mainen

## Abstract

In standard models of perceptual decision-making, noisy sensory evidence is considered to be the primary source of choice errors and the accumulation of evidence needed to overcome this noise gives rise to speed-accuracy tradeoffs. Here, we investigated how the history of recent choices and their outcomes interacts with these processes using a combination of theory and experiment. We found that the speed and accuracy of performance of rats on olfactory decision tasks could be best explained by a Bayesian model that combines reinforcement-based learning with accumulation of uncertain sensory evidence. This model predicted the specific pattern of trial history effects that were found in the data. The results suggest that learning is a critical factor contributing to speed-accuracy tradeoffs in decision-making and that task history effects are not simply biases but rather the signatures of an optimal learning strategy.

## INTRODUCTION

Evidence accumulation is an important core component of perceptual decision-making that allows organisms to mitigate the effects of environmental uncertainty by combining information through time (Roitman and Shadlen, 2002; Chittka *et al.*, 2003; Palmer, Huk and Shadlen, 2005; Gold and Shadlen, 2007; Ratcliff and McKoon, 2008; Bowman, Kording and Gottfried, 2012; Histed, Carvalho and Maunsell, 2012; Brunton, Botvinick and Brody, 2013). Simple theoretical models of evidence accumulation based on a random diffusion-to-bound (DDMs) have been successful in critical aspects of the performance of psychophysical tasks, capturing the dependence of accuracy (psychometric) and reaction time (chronometric) functions. Key elements of these models have begun to be tested both by searching for neural activity corresponding to model variables (Roitman and Shadlen, 2002; Hanks, Ditterich and Shadlen, 2006; Beck *et al.*, 2008; Kiani, Hanks and Shadlen, 2008; Erlich *et al.*, 2015; Hanks *et al.*, 2015) and by the use of more sophisticated task design and modeling (Brunton, Botvinick and Brody, 2013; Zariwala *et al.*, 2013). Yet substantial ambiguity remains concerning nearly all critical features of this class of models, including the basic mechanisms supporting integration, how a bound is determined, and the origins of apparent randomness.

One widely observed but not well-understood phenomenon is that different kinds of decisions appear to benefit from accumulation of evidence over different time scales. For example, monkeys performing integration of random dot motion (Roitman and Shadlen, 2002) and rats performing a click train discrimination task (Brunton, Botvinick and Brody, 2013) can integrate evidence for over one second. But rats performing an odor mixture categorization task fail to benefit from odor sampling beyond 200-300 ms (Uchida and Mainen, 2003; Zariwala *et al.*, 2013). A possible explanation is that neural integration mechanisms that are specific to a given species and sensory modality. However, even animals performing apparently similar odor-based decision tasks can show very different integration time windows (Abraham *et al.*, 2004; Rinberg, Koulakov and Gelperin, 2006). Motivation for speed vs. accuracy, or speed-accuracy tradeoff (SAT) (Khan and Sobel, 2004; Palmer, Huk and Shadlen, 2005; Hanks, Ditterich and Shadlen, 2006), which could change the height of the decision bound, has been proposed as a possible explanation for differences seen across similar studies. However, manipulation of motivational parameters failed to increase the effective integration window in odor categorization, suggesting that other factors must limit decision accuracy (Zariwala *et al.*, 2013).

In DDMs, the chief source of uncertainty is stochasticity in incoming sensory evidence, modeled as Gaussian white noise around the true mean evidence rate (Ratcliff, 1978; Ratcliff and Smith, 2004). It is this rapidly fluctuating noise that accounts for the benefits of temporal integration. The nature and implications of other sources of variability have also been considered in models (Ratcliff, 1978; Ratcliff and Smith, 2004; Ratcliff and McKoon, 2008; Mulder *et al.*, 2012; Brunton, Botvinick and Brody, 2013; Fründ, Wichmann and Macke, 2014), including variability in starting position (Mulder *et al.*, 2012), variability in non-accumulation time (Ratcliff and Smith, 2004), and variability in threshold or bound (Ratcliff, 1978). A potentially important source of variability is trial-by-trial fluctuations in the mean rate of evidence accumulation. Such fluctuations would correspond to uncertainty in the mapping of sensory data onto evidence for a particular choice (Beck *et al.*, 2008; Gold *et al.*, 2008). This mapping could be implemented as the strength of weights between sensory representations into action values (Beck *et al.*, 2008). A combination of weights would then represent a classification boundary between sensory stimuli (Majaj *et al.*, 2015). Weight fluctuations would introduce errors that, unlike rapid fluctuations, could not be mitigated by temporal integration and would therefore curtail the benefits of evidence accumulation (Uchida, Kepecs and Mainen, 2006; Zariwala *et al.*, 2013). Such “boundary” variability (not to be confused with the stopping “bound” in accumulation models) might affect differently particular decision tasks, being particularly important when the mapping from stimulus to action must be learned de novo (Uchida, Kepecs and Mainen, 2006; Zariwala *et al.*, 2013). Indeed, effects of reward-history on choices have been shown experimentally in perceptual tasks (Rescorla and Wagner, 1972; Sutton and Barto, 1998; Busse *et al.*, 2011; Scott *et al.*, 2015).

History effects are considered unwanted biases in psychophysical tasks because each trial is typically constructed to be independent of the preceding trials and stimulus-response rules are fixed. Here, we hypothesized that biases are in fact signatures of an optimal learning strategy that is adapted to natural environments, implying that they only appear “unwanted” (or suboptimal) in situations which those conditions do not hold (Summerfield, Behrens and Koechlin, 2011). Intuitively, an optimal learning agent must always use both priors (history of stimuli, choices and rewards) and current sensory information in proportion to their confidence (Pouget, Drugowitsch and Kepecs, 2016; Drugowitsch and Pouget, 2018). To test this idea formally, we derived the optimal choice policy and learning algorithm for an ideal observer that uses accumulation of evidence and reward statistics to infer choices and updates its stimulus-choice mapping—a Bayesian drift-diffusion model (Drugowitsch and Pouget, 2018).

To test this model, we compared performance in two odor-guided decision tasks that were identical except for the nature of the stimuli. The first was an odor mixture categorization task (Uchida and Mainen, 2003) in which the difficulty was increased by making the stimuli closer to the category boundary, essentially decreasing the contrast between odor categories. We expected that performance in this task would be dominated by uncertainty in the stimulus-choice mapping and therefore benefit less from sensory integration. The second was an odor identification task in which the difficulty was increased by lowering stimulus concentration, in which we expected performance would benefit more from integration. Indeed, the change in reaction times over a given range of accuracy differed markedly between the two tasks, despite being tested in the same animals with all other task variables constant. Standard diffusion-to-bound models could fit performance on either task alone, but not both simultaneously. However, the optimal Bayesian-DDM model could fit both tasks simultaneously and outperformed simpler models without learning and with alternative learning rules. Critically, the introduction of learning predicted a history-dependence of trial-by-trial choice biases whose specific pattern and magnitude were qualitatively and quantitatively matched to the data. These findings suggest that “errors” in many psychophysical tasks are not due to stochastic noise, but rather to suboptimal choices driven by optimal learning algorithms that are being tested outside of the conditions in which they evolved (Beck *et al.*, 2012).

## RESULTS

### Different speed-accuracy tradeoffs in two different olfactory decision tasks

We trained Long Evans rats on two different versions of a two-alternative choice olfactory reaction time task. We refer for convenience to these as two “tasks”, but they were identical in all aspects except for the nature of the presented olfactory stimulus (**Fig.1**). In the first task, “odor identification”, a single pure odor was presented in any given trial and across trials we manipulated difficulty by diluting odors over a range of 3 log steps (1000-fold, liquid dilution) (**Fig.2a**). Thus, the absolute concentration of the odor determined the difficulty. In the second task, “odor categorization”, mixtures of two pure odors were presented at a fixed total concentration but at four different mixture ratios (Uchida and Mainen, 2003) (**Fig.2b**). The distance of the stimulus to the category boundary (50/50, iso-concentration line), determined the difficulty of a given trial, with lower “mixture contrasts” corresponding to more difficult trials. e.g., 56/44 and 44/56 stimuli correspond to 12% mixture contrast were more difficult than 80/20 and 20/80 stimuli, corresponding to 60% mixture contrast. Note that the easiest odor pairs (10^-1^ dilution and 100% contrast) were identical between the two tasks and that both tasks relied on the same partition of the sensory space but applied to different regions of that space. In a given session, the eight stimuli from one of the two tasks were presented in randomly interleaved order. To ensure that any differences in performance were due to the manipulated stimulus parameters, all comparisons were done using the same rats performing the two tasks on different days with all other task variables being held constant (**Fig. S1**).

**Figure 1.**
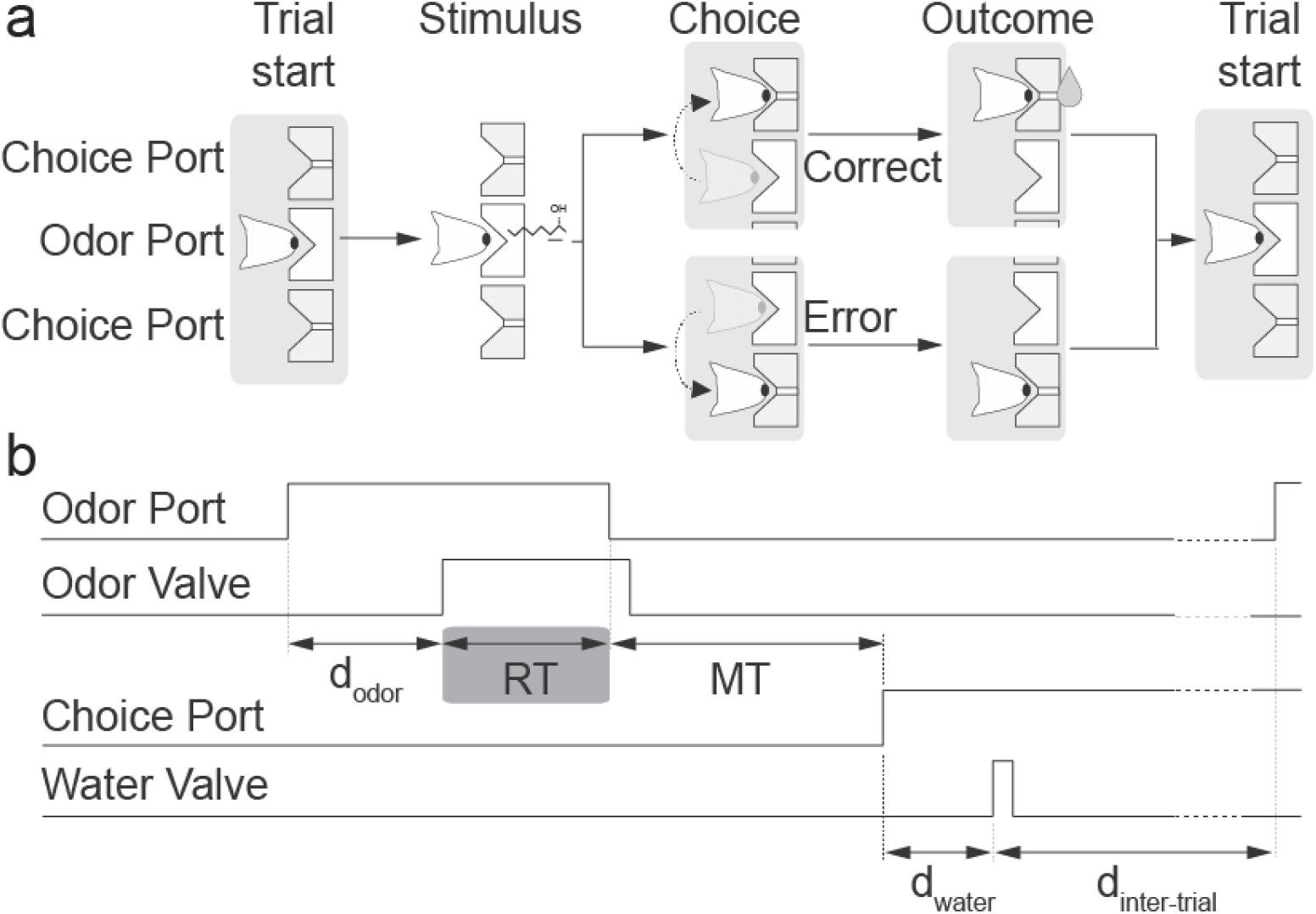
Two-alternative odor choice task. **(a)** Rats were trained in a behavioral box to signal a choice between left and right port after sampling a central odor port. The sequence of events is illustrated using a schematic of the ports and the position of the snout of the rats. **(b)** Illustration of the timing of events in a typical trial. Nose port photodiode and valve command signals are shown (thick lines). A trial is initialized after a rat pokes into a central Odor Port. After a randomized delay *d_odor_* a pure odor or a mixture of odors is presented, dependent of the task at hand. The rat can sample freely and respond by moving into a choice port in order to get a water reward. Each of these ports is associated to one of two odors – odor A ((R)-(–)-2-Octanol) and odor B ((S)-(+)-2-Octanol). Highlighted by the grey box, reaction time (RT) is the amount of time the rats spend in the central Odor Port after odor valve is on (i.e. discounting *d_odor_*). See Experimental Procedures for more details.

**Figure 2.**
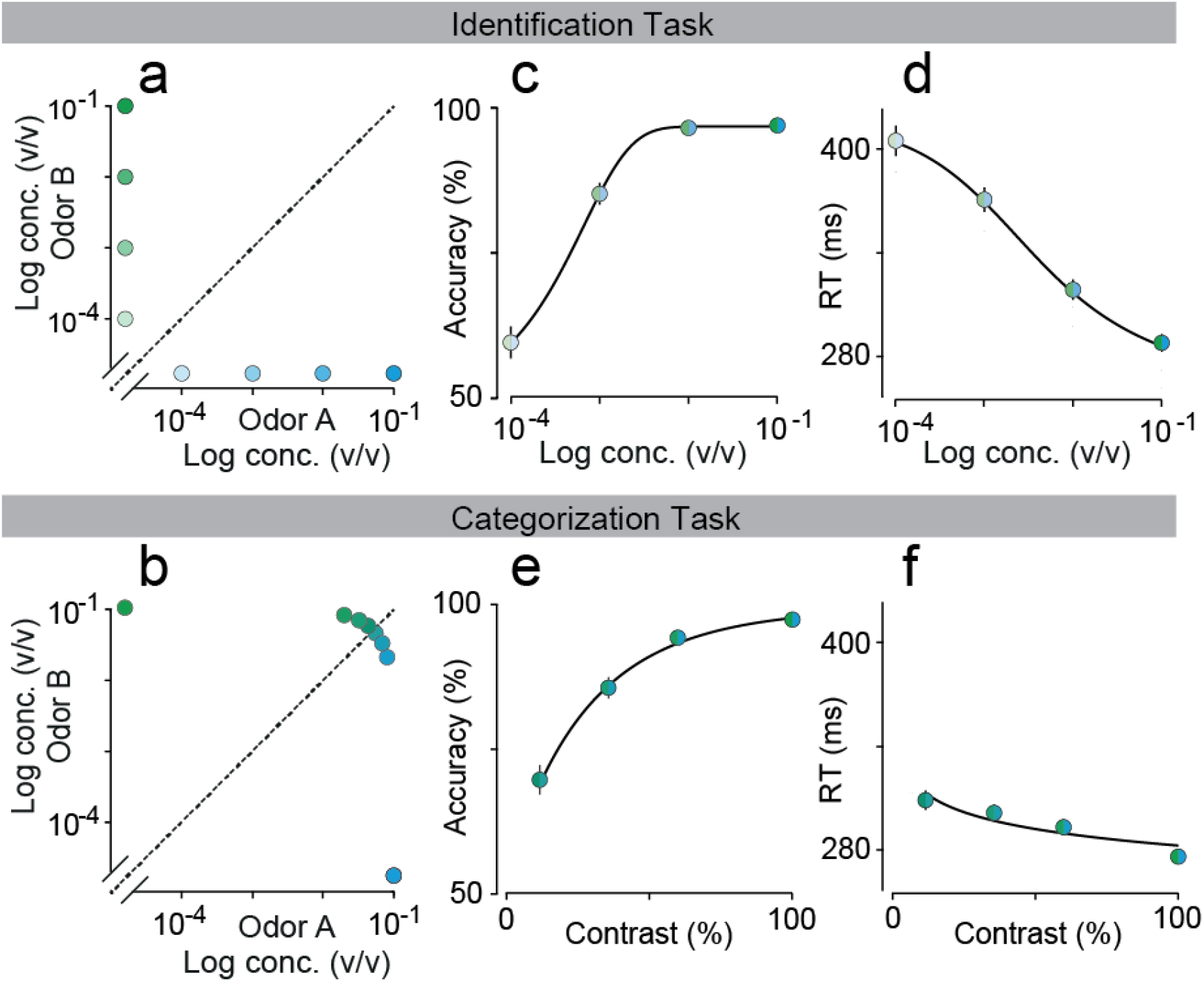
Comparison between odor identification and mixture categorization tasks. **(a,b)** Stimulus design. In the odor identification task, the odorants were presented independently at concentrations ranging from 10^-1^ to 10^-4^ (v/v) and sides rewarded accordingly (a). For the mixture categorization task, the two odorants were mixed in different ratios presented at a fixed total concentration of 10^-1^, and rats were rewarded according to the majority component (b). Each dot represents one of the 8 stimuli presented for each task. **(c,d)** Mean accuracy (c) and mean reaction time (d) for the identification task plotted as a function of odor concentration. **(e,f)** Mean accuracy (e) and mean reaction time (f) for the categorization task plotted as a function of mixture contrast (i.e. the absolute percent difference between the two odors). Error bars are mean ± SEM over trials and rats. Colors in dots are presented as to help parse between stimulus space and psych- and chrono-metric curves. Solid lines depict the obtained fits for the predicted curves of a DDM, an exponential curve for performance and a hyperbolic tangent for RTs, as described in Palmer, Huk and Shadlen, 2005.

We quantified performance using accuracy (fraction of correct trials) and odor sampling duration, a measure for reaction time (RT) (Uchida and Mainen, 2003; Zariwala *et al.*, 2013) (**Fig.1, Fig. S2** for individual rats). We observed that rats performing the two tasks showed marked differences in how much reaction times increased as task difficulty was increased (**Fig. 2c-f**). For the identification task, reaction times increased substantially (112 ± 3 ms; mean ± SEM, n = 4 rats; F(3,31) = 44.04, P < 10^-7^; **Fig. 2d**), whereas for the same animals performing the categorization task, the change was much smaller (31 ± 3 ms; F(3,31) = 2.61, P = 0.09, ANOVA) (**Fig. 2f**), despite the fact that the accuracy range was similar.

In order to control for the possibility that a slightly smaller range of performance accuracy for the categorization task accounted for differences in SAT, we re-ran this task with two sets of stimuli with wider ranges of mixture contrasts including harder, lower contrast stimuli. This yielded a range of accuracies as broad as those in the identification task (**Fig. S3a**). The change in RTs across all difficulties was 41 ± 24 ms and 50 ± 19 ms for the two datasets tested (**Fig. S3b**), slightly higher than the one observed for the original categorization dataset, yet still much smaller than the RT change for identification (**Fig. S3c, d**). Therefore, the difference observed in SAT for odor identification vs. mixture categorization was not due to differences in the range of task difficulties.

### Construction of a diffusion-to-bound model for olfactory decisions

In order to explore further which variables might be constraining the rats’ behavioral performance, we fit the data using DDMs (DDM, **Fig. 4a**). In two alternative forced choice tasks, in which the subjects have the freedom to respond at any time within a trial, trading off the cost associated with accumulating evidence for slower, more accurate choices with the lower expected reward for faster, less accurate choices becomes paramount. With adequately tuned decision boundaries, DDMs are known to implement the optimal, i.e., reward-rate maximizing, strategy to this tradeoff in a wide range of different tasks, including the ones used in the present study (Bogacz *et al.*, 2006; Drugowitsch *et al.*, 2012; Tajima, Drugowitsch and Pouget, 2016).

Here we implemented a DDM composed of sensory, integration and decision layers. The sensory layer implements a transformation of concentrations into momentary evidence for odors A and B. Perceptual intensity in olfaction (Stevens, 1975; Wojcik and Sirotin, 2015), as in other modalities (Palmer, Huk and Shadlen, 2005; Brunton, Botvinick and Brody, 2013) can be well-described using a power law. We therefore defined the mean strength of sensory evidence *μ* for each odor using a power law of the odor concentration (**Fig. 2a-b**),

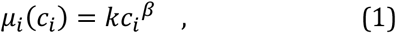

where k and *β* are free parameters (Palmer, Huk and Shadlen, 2005). We constrained k and *β* to be identical between the two odors, which were stereoisomers with identical vapor pressures and similar intensities (Taniguchi, Kashiwayanagi and Kurihara, 1992; Pierce *et al.*, 1995; Uchida and Mainen, 2003; Laska *et al.*, 2004). Evidence at each time step is drawn from a normal distribution *m_i_*(*t*): *N*(*μ_i_, σ*), where *σ* = 1 is the standard deviation of the variability corrupting the true rate, *μ_i_*. The integration layer, which also consists of two units, integrates the noisy evidence over time independently for each odor. The last step of the model consists of a unit that takes the difference between the two integrated inputs. If this difference exceeds a given bound, *θ* or –*θ*, the model stops and makes a choice according to the bound that was hit: left for *θ*, right for –*θ*. Finally, we allowed for a time-dependent linear decrease in the bound height (“collapsing bound”), *τ*, mimicking an urgency signal (Churchland *et al.*, 2011; Drugowitsch *et al.*, 2012) (see Experimental Procedures for details).

### Diffusion-to-bound model fails to fit both tasks simultaneously

In an attempt to explain our behavioral data with the standard DDM, we developed a series of different fitting procedures in order to test how well this model describes the data. These fitting procedures all involved maximizing a log-likelihood function for a data set of 22,208 (identification), 19,947 (categorization) or 42,155 (both) trials using simulations over 100,000 trials (see Experimental Procedures). The overall quality of the fit obtained with each procedure is shown in **Fig. S4**. The first approach we considered was to test whether we could predict the behavioral data of one task using the fitted parameters from the other task. The free parameters of the standard DDM were first fitted to behavioral data of the identification task. The model captured the overall behavior of the rats in this task. As model performance dropped from 93.7% to 56.4% (data: 97.0 ± 0.9% to 59.4 ± 2.7%; mean ± SEM, n = 22,126 trials/4 rats) with decreasing odor concentration, mean reaction times increased from 286 to 401 ms (data: 290 ± 2 to 402 ± 3 ms) (**Fig.4b**, black line). This is because the evidence for lower concentrations is dominated by noise, making the signal to noise-ratio smaller. The model therefore takes longer to reach a bound while being more prone to hitting the wrong bound first.

Next, we asked whether we could predict the rats’ psycho- and chronometric curves in the categorization task using the model with the parameters we had fitted to the identification task. As shown in **Fig. 4c**, the model had reaction times within the same range; the model’s reaction time increased from 285 to 311 ms (data: 279 ± 1 to 311 ± 2 ms; mean ± SEM, n = 19,924 trials/4 rats). But the model strongly overestimated the animals’ accuracy at low odor contrast (e.g. model 93.4% vs. data 69.0 ± 1.3% at 12% mixture contrast) (**Fig. 4c**, black line). As a second procedure, we attempted to fit the model parameters to the categorization task and to predict the identification task (**Fig. 4b-c**, dashed lines). This was also unsuccessful: the model predicted much faster (340 ms) responses than what is seen in the data for lower concentrations. A third procedure, simultaneous fits, also failed in describing both tasks successfully (**Fig. S5**). The only satisfactory description of our data for this model was to consider both tasks independently (**Fig. 4b-c**).

### Differences in SAT are not due to context dependent strategies

Motivational variables can modulate performance and reaction time in perceptual tasks. For example, variables like reward rate (Drugowitsch *et al.*, 2012) or emphasis for accuracy vs. speed (Palmer, Huk and Shadlen, 2005; Hanks, Ditterich and Shadlen, 2006) can have an effect on observed SATs, by modulating decision criteria. This could result in changes in non-stimulus dependent parameters such as integration threshold *θ,* non-decision time *t_ND_* and lapse rate from one task to the other. Because identification and categorization tasks were run in separate sessions, we also considered the possibility that rats might have shifted their decision criteria between tasks. To address this, and to cover the stimulus space more thoroughly, we devised a “mixture identification” task in which we interleaved the full set of stimuli from the categorization and identification tasks as well as intermediate mixtures. Thus, on each trial the stimulus was chosen randomly from one of four mixture ratios at one of four concentrations (**Fig. 3a**). Consistent with the previous observations, RTs in this joint task were strongly affected by concentration but considerably less so by mixture contrast (**Figs. 3b-c**). A two-way ANOVA showed that RTs changed significantly across the different odorant concentrations (F(3,48) = 8.69, *P* < 10^-3^); but for a given total concentration of the odorants, this change was not significant across the different mixture contrasts and subjects (F(3,48) = 0.94, P = 0.42). There was no significant interaction of odorant concentration and mixture contrast (F(9,48) = 0.28, P > 0.9). For individual subjects, all 4 rats showed significant effect of odorant concentration (ANOVA for each rat: F_1_(3,15)=78.66, P_1_<10-^6^; F_2_(3,15)=14.66, P_2_<10-^3^; F_3_(3,15)=204.91, P_3_<10^-7^; F_4_(3,15)=27.86, P_4_<10^-4^), while only 2 showed significant modulation of mixture contrast across sessions (F_1_(3,15)=1.14, P_1_=0.39; F_2_(3,15)=0.52, P_2_=0.67; F_3_(3,15)=9.6, P_3_<0.01; F_4_(3,15)=6.47, P_4_<0.05). Additionally, the best fitting DDM model could not explain this data-set successfully (**Fig. S6**). These results indicate that the differences in the relation between accuracy and reaction time for the two tasks are not due to differences in decision criteria.

**Figure 3.**
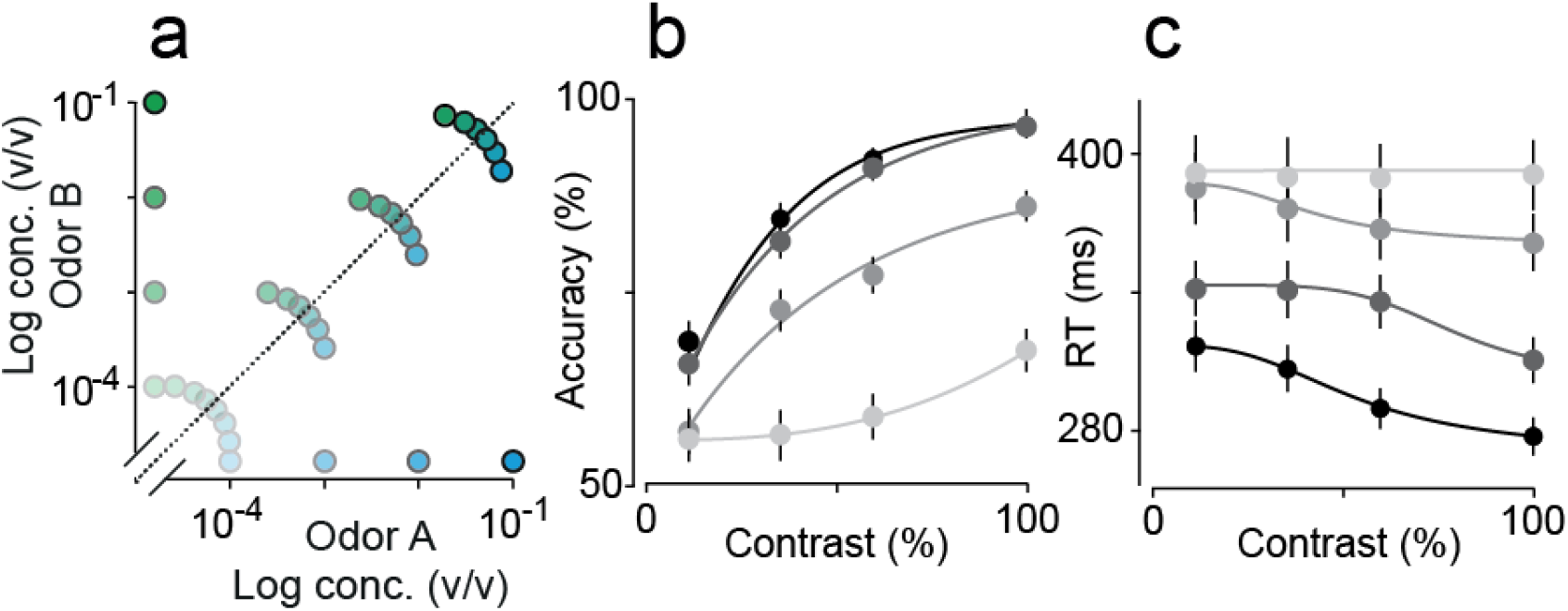
Odor mixture identification task. **(a)** Stimulus design. Two odorants (S-(+)-2-octanol and R-(-)-2-octanol) were presented at different concentrations and in different ratios as indicated by dot positions. In each session, four different mixture pairs (i.e. a mixture of specific ratio and concentration and its complementary ratio) were pseudo-randomly selected from the total set of 16 mixture pairs and presented in an interleaved fashion. **(b, c)** Mean accuracy (b) and mean of reaction times (c) plotted as a function of mixture contrast. Each point represents a single mixture ratio. Error bars are mean ± SEM over trials and rats. Solid lines depict the obtained fits for the predicted curves of a DDM for each set of mixtures at a particular concentration, family functions as described in Palmer, Huk and Shadlen, 2005. Colors represent the total concentration of the mixture, with black indicating a 10^-1^ mixture and lightest grey 10^-4^ mixtures.

Similar results were obtained for other variants of the integration-to-bound model, such as variants of the accumulator model of two-choice discrimination (Smith and Vickers, 1988) and the two-race competition model (Usher and McClelland, 2001) (not shown). These results suggest that the data are not compatible with the standard assumptions of this class of models.

### Diffusion-to-bound model with stimulus dependent Bayesian learning fits performance across both tasks

Until now we have been considering that all behavioral uncertainty comes from moment-by-moment stochastic fluctuations in incoming sensory evidence. Although it is standard in psychophysical tasks to assume that previous trials’ choices and outcomes do not bias current responses and that each trial is independent of all others, it is well known from reward-based decision-making tasks that subjects’ choices are sensitive to the recent history of rewards (Herrnstein, 1974; Baum, 1979; Sugrue, Corrado and Newsome, 2004). Indeed, it has been shown that rodents exhibit trial-history dependent biases in psychophysical tasks (Busse *et al.*, 2011). One possible explanation for the overestimate of accuracy in the categorization task is that these trial-history effects impact this task preferentially (Uchida, Kepecs and Mainen, 2006; Zariwala *et al.*, 2013). Reward expectation, for instance, has been shown to influence performance and RTs in perceptual tasks (Lauwereyns *et al.*, 2002; Roesch and Olson, 2004; Zariwala *et al.*, 2013), implying that perception can actually be influenced by rewards.

Taking this knowledge into consideration, we hypothesized that on-going fluctuations in the animals’ mapping from odors to choice directions or, equivalently, their representation of the categorical boundary, might limit performance. Such fluctuations might be generated by reward expectations and reinforcement-driven learning processes. To explore this, we decided to take a look at performance and choice biases (Busse *et al.*, 2011).

To test this idea, we asked how rodents could use rewards to update their category “boundary”, i.e, the line that divides stimuli associated with right and left choices (see **Fig. 2a,b**). To do so, we assumed a volatile world that requires a continuous updating of this category boundary and used Bayesian decision theory to derive the near-optimal updating strategy under these circumstances. This strategy is close-to-optimal in the sense that it well-approximates the best possible category boundary estimate given all available information, and under the assumption that the “true” category boundary drifts stochastically across consecutive trials (see Experimental Procedures for details). Even though it approximates the optimal strategy, which is intractable, it yields behavioral performance indistinguishable from optimal (Drugowitsch and Pouget, 2018). The use of this strategy resulted in a diffusion-to-bound model with stimulus-dependent Bayesian learning (which we refer to as “Bayes-DDM”) (**Fig. 5a**). The Bayes-DDM is augmented with weights that transform the stimulus input into evidence:

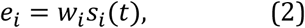

which is then combined with bias *b* to form a net evidence

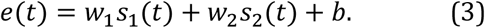

In this equation the weights *w_i_* and the bias *b* define, respectively, the slope and offset of the category boundary.

After each trial we updated the stimulus weights *w_i_* using a tractable approximation to the intractable Bayes-optimal learning rule,

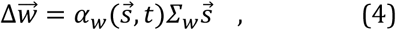

where *Σ_w_* is the weight covariance matrix (also learned; see Experimental Procedures), that quantifies the current weight uncertainty, and 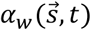 the learning rate. This learning rule introduces three new parameters that describe the learner’s assumptions about how the weights change and influence the learning rate α (see Experimental Procedures for details).

We tested the ability of Bayes-DDM to capture performance in both olfactory decision tasks. We fit this model through log-likelihood maximization of both tasks simultaneously (Palmer, Huk and Shadlen, 2005). However, as the model was adjusting its weight and bias parameters across consecutive trials, and was using collapsing boundaries, we could not apply the same or even any closed form analytical predictions (Ratcliff, 1978). Thus, we found the predicted mean RTs, choice probability and trial-to-trial choice biases by numerically simulating a sequence of 100,0 trials for any combination of parameters. These were then fit to a data set of 22,208 (identification) and 19,947 (categorization) trials by log-likelihood maximization (see Experimental Procedures). The inclusion of the new category boundary learning parameters in the DDM allowed the simultaneously fit of the two tasks (**Fig. 5b-c**). The Bayes-DDM showed a decrease of performance from 97% to 65%, accompanied by an increase of 29 ms in reaction time for the categorization task and a deterioration of performance from 96% to 62%, with a 93 ms increase in reaction time for the identification task. The model showed similar results as to what was observed in the behavioral data (**Fig. 5**).

As a further test we assessed whether the model could fit the behavioral results for all the intermediate concentration levels of the mixture task (**Fig. 3 b-c**). To do so, we fitted the model to the 32 stimuli from the interleaved condition (**Fig. 3a**). We found that the model was indeed able to fit well the full set of psycho- and chono-metric functions (**Fig. 5d**, solid lines).

### Bayes-DDM successfully predicts trial-by-trial conditional changes in choice bias

The Bayes-DDM model can be considered as a hypothesis concerning the form of trial-to-trial biases that we expect to be sufficient to explain the data. Crucially, the specific predictions of this model can be tested against behavioral variables that were not directly fit. That is, we can check whether the form of the trial-to-trial biases in the experimental data is in fact compatible with the form and magnitude of the learning we introduced.

To do so, we first quantified the impact of a previous trial by calculating the difference in the average choice bias conditional upon the trial being correct *and* a given stimulus being delivered (**Fig.6a,b**), relative to the overall average choice bias (highlighted by the red arrows in **Fig. 6a**) (see Experimental Procedures). Because Δ*C_B_(x)* was symmetric for left/right stimuli (**Fig. 6a, b**; **Fig. S7**), we plot *ΔC_B_*(*x*) collapsed over stimuli of equal difficulty (**Fig. 6c,d; Fig. S8**). Note that *ΔC_B_*(*x*) measures the fractional change in choice probability, with *ΔC_B_*(*x*) > 0 indicating a greater likelihood of repeating a choice in the same direction as the prior trial, *ΔC_B_*(*x*) < 0 indicating a greater likelihood of making a choice in the opposite direction. We also calculated the equivalent updating curves conditional on the trial being *incorrect* and a given stimulus being delivered. However, likely due to the much smaller number of error trials (rats are on average 85% correct for all tasks), especially for easy trials, there was a great deal more variability in these curves (**Fig. S9**). As a result, our data did not sufficiently constrain the fits to these updating curves, particularly for the categorization task (**Fig. S10**). We therefore focused on the analysis of the dependence of correct trials for the two tasks.

These analyses showed that rats have a tendency to repeat a choice in the same direction that was rewarded in the previous trial (“win-stay”; **Fig. 6c,d**). But the stimulus-dependent analysis revealed a qualitative difference between the identification and categorization tasks with respect to how the stimulus in the past trial impacted the change in bias (**Fig. 6c, d**). For the identification task, the influence of the previous trial was largely stimulus independent (**Fig. 6c**, one-way ANOVA, F(3,12) = 2.0, *P* = 0.17). For the categorization task (**Fig. 6d**), in contrast, the influence of the previous trial showed a graded dependence on the stimulus, being larger for a difficult previous choice than for an easier one (F(3,12)=25.4, P<10^-5^).

The Bayes-DDM generated predictions for the shape and amplitude of history-dependent choice bias (updating) functions for both tasks (lines in **Fig. 6c and 6d**). Importantly, these functions were predicted rather than directly fit, since the only data used for the fits were the trial-averaged accuracy and RT curves. An important feature of the Bayes-DDM is that it depends on both the accumulated inputs 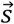 and decision time ř, reflecting a form of decision confidence (Drugowitsch and Pouget, 2018; see Experimental Procedures). In tasks like ours, with a varying difficulty, harder trials are associated with later choices and come with a lower decision confidence (Kiani and Shadlen, 2009). On correct trials, the learning rate is smaller when the animal’s confidence is high. This makes sense: if the animal is correct and highly confident, there is little reason to adjust the weights. However, continual bias learning washes out the confidence – reaction time relationship, such that neither data no the Bayes-DDM model feature a strong modulation of learning by reaction time (**Fig. S11**).

Remarkably, for both tasks the predictions of the Bayes DDM model closely matched the data. For the categorization task, as expected, the model captured the strong dependence of the updating curve on stimulus difficulty (**Fig. 6d**). This is due to the fact that for easy stimuli, the predicted probability of a rewarded trial (i.e. the value of the evidence at stopping time) is nearly equal to 1 and there is little surprise (i.e. high confidence) and little learning. For the identification task the model was also able to capture the relative lack of stimulus dependence of the updating curve (**Fig. 6c**). This is explained by the fact that in the identification task the signal-to-noise ratio is low for the most difficult trials, implying that the sensory component of Eq. 3 will be low. Thus, there is a larger contribution of the stimulus-independent updating term (the bias – b) in updating choice bias (see Experimental Procedures).

### Comparison with other learning rules

The optimal Bayes DDM learning rule takes a complex form involving multiples terms who respective roles are not immediately clear. In order to gain some insight into why this rule captures the animals’ behavior, and whether confidence plays a role, we fitted several variations of our model and used Bayesian model comparison to determine which one best accounted for our data.

We first fitted a model without learning but in which the weights are drawn on every trial from a multivariate Gaussian distribution whose mean is set to the optimal weights (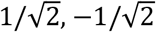 and 0) and whose variance is a free parameter. Interestingly, this model could fit the psychometric and chronometric curves in both tasks with a single DDM. However, the model failed to show sequential effects in either task, as one would expect since the weights are redrawn independently on every trial (**Fig. S12**). Bayesian model comparison confirmed that this model performs considerably worse than the one using the optimal learning rule (**Fig. 7**).

Next, we tried a model with a limited form of learning in which the optimal Bayes DDM learning rule is applied only to the bias while the sensory weights are set to their optimal values. It has indeed been recently argued that sequential effects can be captured by variations in the bias (Busse *et al.*, 2011). This model has a BIC score comparable to the optimal model (**Fig. 7**) and captures the flat profile of the sequential effects in the identification task, thus suggesting that sequential effects in this task are due to fluctuations in the bias. However, this model fails to account for the profile of sequential effect in the categorization task (**Fig. S13**). While the data shows sequential effects inversely proportional to the difficulty of the previous trial, the model predicts a flat profile, similar to what we observed for the identification task (**Fig. S13g**).

Conversely, we fitted a model in which the sensory weights, but not the bias, are adjusted on every trial according to the optimal learning rule. This model fits to the psychometric and chronometric curves reasonably well in both tasks (**Fig. S14**). However, in contrast to the previous model, this one captures the sequential effects in the categorization task but not in the identification task (**Fig. S14d**). Moreover, the BIC score for this model is far worse than for the optimal model. These last two models combined reveal that the learning-induced fluctuations in the bias is what allows the optimal model to capture the sequential effects in the identification task, while the learning-induced fluctuations in the weights play a critical role in capturing the sequential effects in the categorization task.

Finally, we explored a possible implementation of the Bayes-DDM rule by considering a simple delta rule that is modulated by decision confidence. We aimed to develop a last heuristic model: a neural network that mimics our Bayes-DDM. For this purpose, we combined a standard DDM with a delta rule of the form:

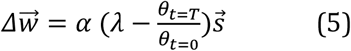

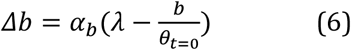

where *θ_t=0_* is the value of bound at the beginning of the trial while *θ_t=τ_* is the value of the bound at the time of the decision, *T, λ* is the correct choice (1 or -1), and *a* and *a_b_* are the weight and bias learning rates. The modulation of learning by confidence is due to the term 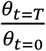. The collapsing bound causes this ratio to decrease with the time elapsed since the beginning of the trial. Critically, elapsed time is inversely proportional to confidence in DDM when the difficulty of the task is unknown and varied from trial to trial (Kiani and Shadlen, 2009; Drugowitsch *et al.*, 2012), as is the case in our tasks. Therefore, the error term in this learning rule is decreasing as confidence decreases over time, which is to say the model learns less when it is less confident, which makes intuitive sense. Ultimately, however, this rule is only an approximation to the optimal rule. Nonetheless, Bayesian model comparison revealed that this learning rule accounts for our experimental data about as well as the full optimal learning rule, thus indicating that the rat’s behavior is consistent with a confidence weighted learning rule (**Fig 7, Fig. S16, RL-DDM**).

The results of Bayesian model comparisons are often sensitive to the way extra parameters are being penalized. Importantly, we found this not to be the case in our data as the ranking of the models remains the same whether we use AIC, AICc or BIC. Moreover, our conclusions hold whether we fitted individual animals separately, or fitted all the data at once as if it was obtained from a single ‘meta-rat’ (**Fig. 6, Fig. 7** and **Fig. S4**).

### Fluctuations in category boundary degrade odor categorization performance more than identification

Finally, we sought to use the Bayes-DDM model to gain insight into how category boundary learning works in conjunction with integration-to-bound to explain the difference between identification and categorization task performance. To do so, we analyzed the dynamics of the weights in relationship to sensory evidence (**Fig. 8**). For each simulated trial, we considered the accumulated evidence for each side and divided it by the total integration time; we termed this the “inferred drift rate” of a trial (see Experimental Procedures). First, we plotted the simulated trials for the standard DDM that was fit to the identification task (**Fig. 8a-b**). Remember that this version of the model generated overly high accuracy in categorizing mixtures (solid black lines from **Fig. 4**). Here, each dot is a simulated trial and the scatter of dots around each stimulus reflects the impact of stochastic noise in the DDM. In this representation, the ideal category boundary is a line with slope equal to 1 with all stimuli below this line categorized as “left” choices and all stimuli above this line as “right” choices. In **Figure 8c, d**, we plot one of the most difficult stimuli for each task (correct choices in blue and errors in red). For the ideal category boundary, it can be seen that performance on the categorization task is expected to be much higher than for the identification task.

**Figure 4.**
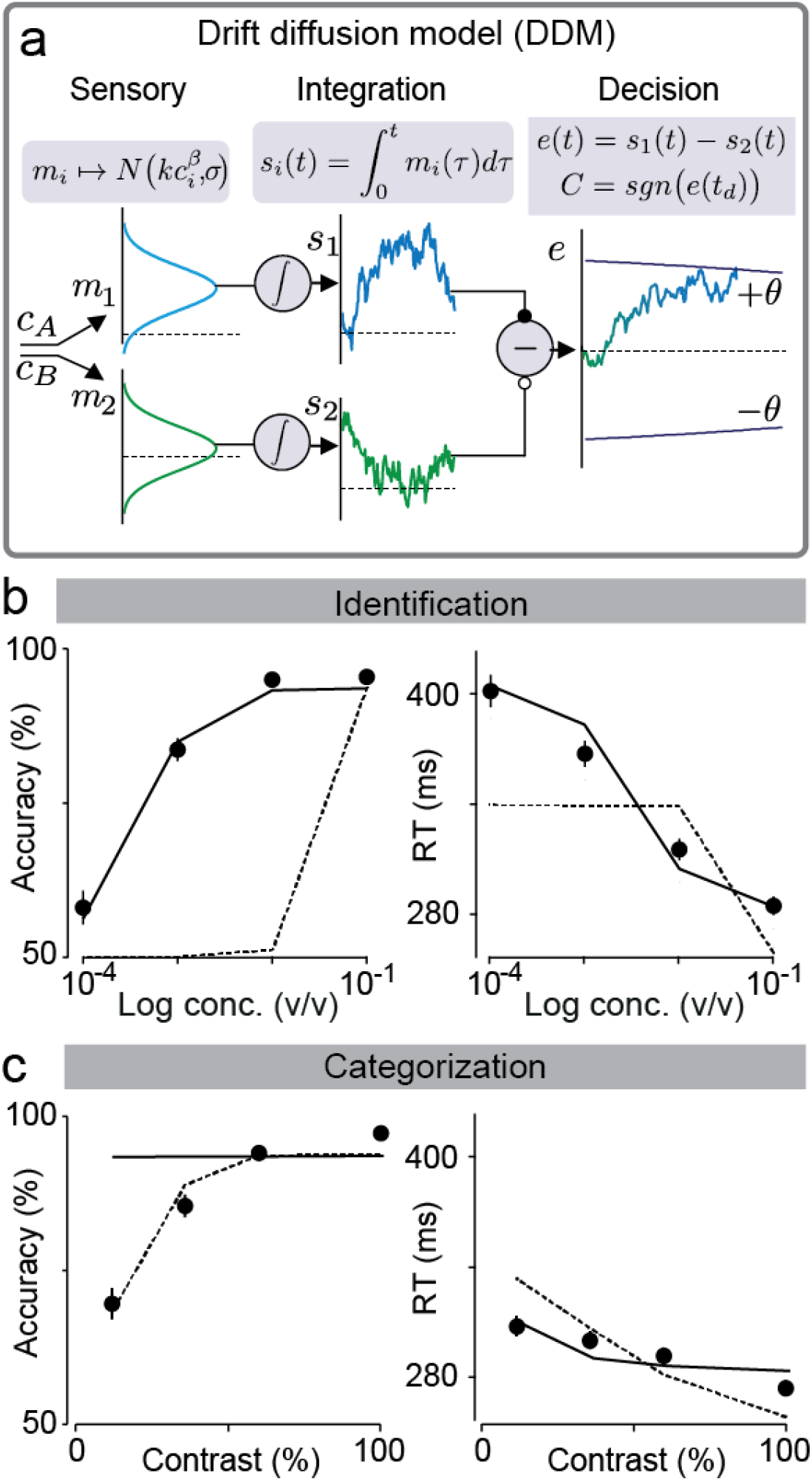
Failure to simultaneously fit performance on identification and categorization tasks with Drift-diffusion model. **(a)** Drift-diffusion model (DDM). The model consists of three layers – Sensory, Integration and Decision layer. At the sensory layer, concentrations are transformed into rates that are contaminated with Gaussian noise. These rates are then integrated over time (Integration layer) and combined. Note that the choice of weights (-1 and 1) for the Decision layer allows it to effectively be a Drift-Diffusion model with collapsing bounds. This model presents 7 parameters (see Experimental Procedures). **(b)** Fitting results for accuracy and reaction time in identification task. Black solid line represents the model fit for this data, and dashed lines the prediction from the categorization data fit. **(c)** Fitting results for accuracy and reaction time in categorization task. Solid black lines depict the prediction for this data from the model fitted to identification, and dashed lines the DDM fit for this data. Error bars are mean ± SEM over trials and rats.

In **Figure 8e, f** we show the effect of learning, which introduces variability in the category boundary in the Bayes-DDM (grey area indicates 1 standard deviation from the mean). The plot shows that for the identification task, learning-induced boundary fluctuations changed the classification of very few trials while, in the categorization task, many were affected (grayed points unchanged, red/blue points changed).

**Figure 5.**
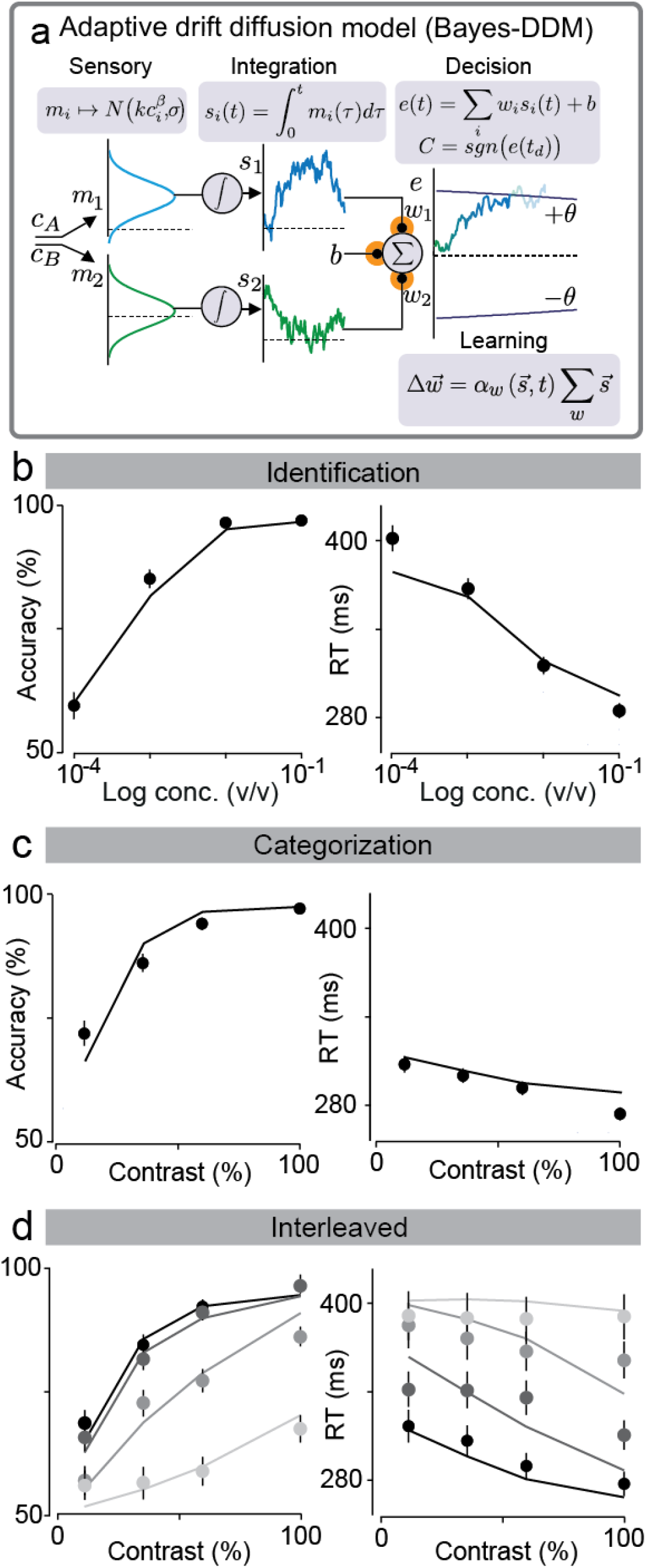
Bayesian DDM with bias and stimulus learning explains identification and categorization task simultaneously. **(a)** DDM is even further expanded with the addition of changing stimulus weights, *wı* and *w2,* and trial-by-trial reward dependent bias b. These weights are then combined with the integrated momentary evidences (*s_1_, s_2_*) plus the offset set by the bias b. After each trial the model updates stimulus weights according to the obtained outcome through a Bayesian learning rule. This model has 10 parameters (see Experimental Procedures). **(b,c)** Choice accuracy (fraction of correct trials) and odor sampling duration in identification task (b) and categorization task (c). Solid black line represents model fitted to both tasks (see Experimental Procedures for more details). **(d)** Choice accuracy and odor sampling duration for Interleaved condition. Solid lines represent the obtained fits for Bayes-DDM to this particular data, going from lowest total concentration (lightest grey) to highest (black). Error bars are mean ± SEM over trials and rats.

**Figure 6.**
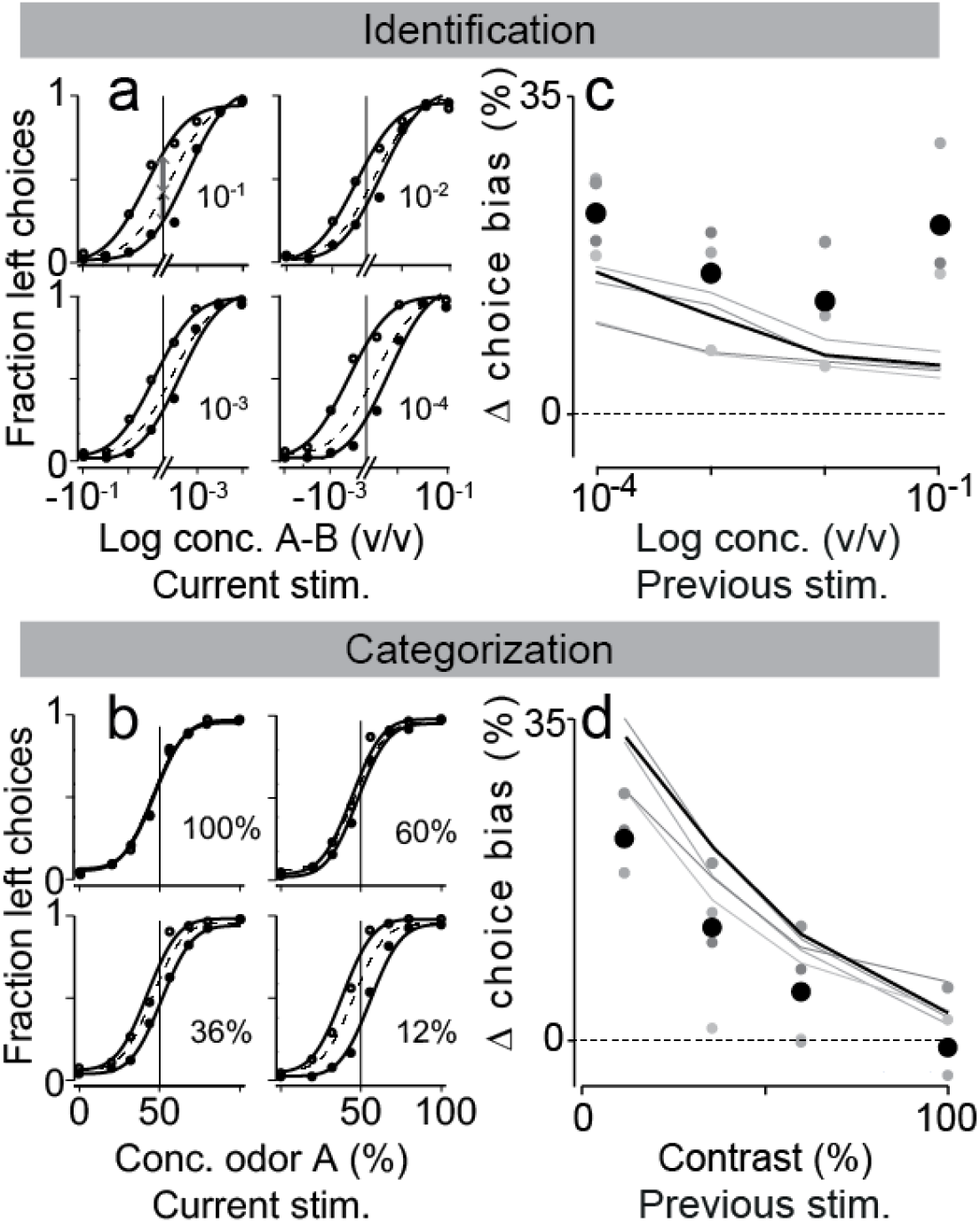
Bayes-DDM predicts trial-by-trial change of choice bias after a rewarded trial and given stimulus. **(a,b)** Psychometric functions for mean (dashed line) and conditional curves following a reward, side choice and a given stimuli. For the identification task all four stimuli from 10^-1^ to 10^-4^ are depicted (a). For the categorization task from 12 to 100% mixture contrast (b). Filled-in circles and equivalent lines represent trials following a right-reward (B>A) and open circles, trials following a left-reward (A>B). Lines represent fits of a cumulative Gaussian to the data. Grey arrows in (a) elucidate the measured change in choice bias (see Experimental Procedures). **(c,d)** Change in choice bias plotted as a function of the previous stimuli, for plots in (a,b). All four different odor concentrations for identification task (c); and all mixture sets for categorization (d). Points correspond to behavioral data (left side), and solid lines to the predicted change from the model fitted to **Fig. 5** (right side). Black points and line correspond to the obtained measurements considering all data together and predicted size of effect when Bayes-DDM fits chrono- and psycho-metric curves for both tasks. Depicted is also each one of the four rats as points with a different shades of grey, with their analogous model predictions (grey lines).

**Figure 7.**
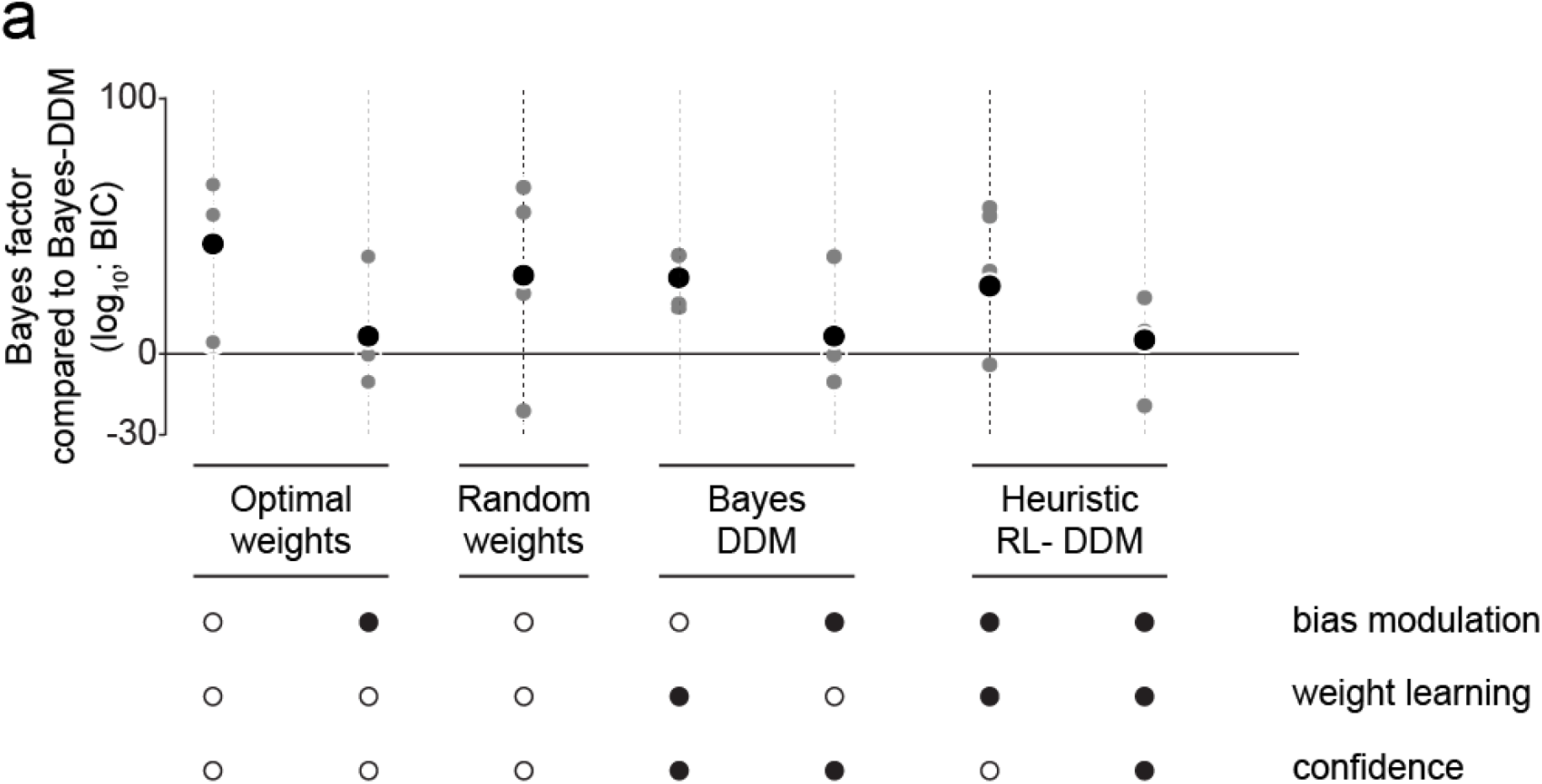
Bayesian model comparison. **(a)**. Model comparison between different family of models against optimal model (Bayes-DDM) using Bayes factor. While the models were only fitted on the mean psycho- and chronometric function, the quality of the fitted data was evaluated considering an objective function additionally takes each model’s prediction of the conditional psychometric functions regarding previous reward into account (see Experimental Procedures for details). Here we depict: two optimal DDM that don’t learn weights, one with trial-by-trial bias modulation and one without; a random weights model in which stimulus-to-bound weights fluctuate from trial to trial stochastically; two simplified versions of Bayes-DDM: one without bias modulation and another without weight learning; and two heuristic models in which we implement a network that mimics a DDM that learns the stimulus-to-choice map through Reinforcement learning (RL-DDM), being one of them without confidence weighted learning. The grey dots show the Bayes factor for individual rats, and the black dots the Bayes factors for the fits using the accumulated data across all rats (adjusted for the increased number of trials). For more details about these models and other versions see Experimental Procedures and Supplementary material.

**Figure 8.**
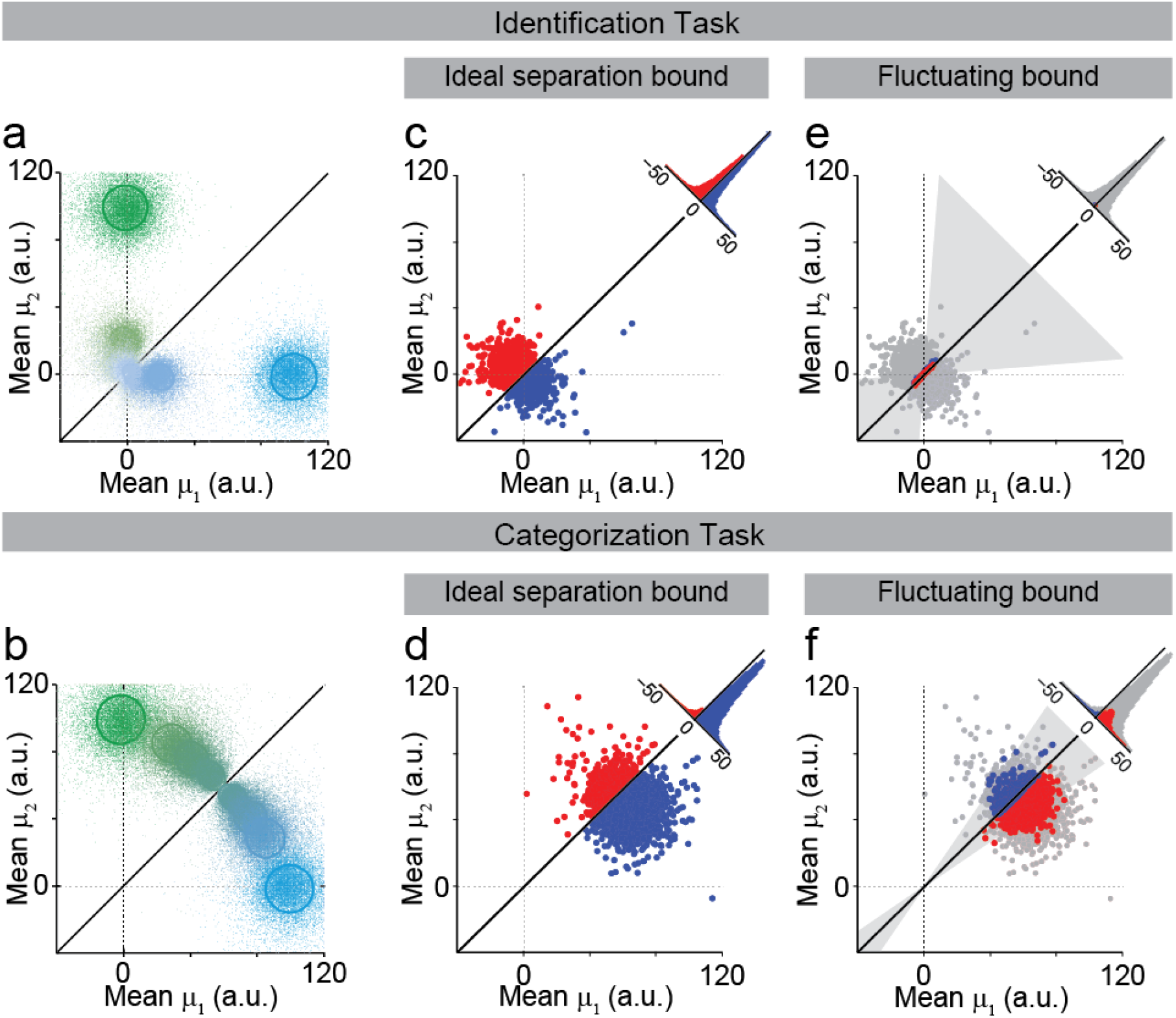
Weight fluctuations amplify errors in categorization task. **(a,b)**. Stimulus space for categorization task. Each point represents a combination of inferred drift rates for a given trial in the pure DDM (with no-learning) that was fit to identification but outperformed in the categorization task (see Experimental Procedures for more details). Solid oval lines represent the Mahalanobis distance of 1 in relation to the population average for each of the eight stimuli. Solid black line depicts the ideal classifying process: above it implies a right-side decision, below it a left. Color code for each point and line follows the same logic as **Fig. 2a,b.** The larger overlap of each set in the identification task (a) explains the performance degradation, as most points are located around the origin. For the categorization task (b), the lack of overlap between stimuli clarifies the higher performance seen in **Fig. 4. (c,d)** Mean drift rates for the most difficult left-decision choice for Bayes-DDM model. In the case of the identification task this is the 10^-4^/0 stimuli (c); for the categorization task the stimulus is the 56%/44% mixture (d). Blue signal the correct classified choices and red the incorrect. Considering the ideal separation bound we show the projected histograms for the difference. **(e,f)** Same as (c,d) but now with fluctuating weights depicted as the slope of the category bound. Grey area indicates the weight fluctuations that are equivalent to 1 standard deviation for both tasks. Blue indicates trials that were originally incorrect in (c,d) but became correct, and red indicates trials that became incorrect but originally correct. Light grey dots indicate answers that remained unchanged. Histograms quantify the four populations of dots.

The difference in effects can be understood by considering that in the Bayes-DDM stimulus weights have a multiplicative effect on evidence strength. Thus, stimulus weight fluctuations correspond to rotations around the origin; the effects are larger for larger stimulus values. Therefore, the difficult mixture stimuli of the categorization task, which have higher concentrations, are much more susceptible to these fluctuations. In the identification task, on the other hand, because stimuli are low concentration and around the origin these changes in the slope of the categorization boundary have little to no effect (**Fig. 8e**). Conversely, we also analyzed the effects of bias in both tasks. In this stimulus representation, changes in *b* affect the intercept of the bound (See Experimental Procedures). However, for bias the effect is similar in both tasks, suggesting that most new errors for mixture categorization originate from fluctuations in the slope of the category bound (**Fig. S17**).

## DISCUSSION

Our results demonstrate that rats show different speed-accuracy tradeoffs depending on the task at hand. When challenged to identify odors at low concentrations, rats show a significant increase of reaction time that is accompanied by performance degradation (**Fig. 2c,d**). In contrast, when the challenge is to categorize mixtures of two odors in different proportions, rats show only a small increase in reaction time (**Fig. 2e,f**). We used a standard drift-diffusion model (DDM) to show that this task difference cannot be explained by stimulus noise (**Fig. 4**) even with the addition of reward-dependent choice biases (**Fig. S13**). We therefore introduced a Bayesian learning process, the kind theorized to drive stimulus-response learning (Bayes-DDM) optimally in dynamic environments, that is, environments in which the optimal weights evolve over time (Drugowitsch and Pouget, 2018). With the combination of these three factors – stimulus noise, reward bias and categorical boundary learning – the Bayes-DDM model not only fit the average performance data (**Fig. 6b-e**), but also predicted the choice biases on the recent history of stimuli, choices and rewards (**Fig. 6f,g**). Furthermore, the Bayes-DDM model was able to fit the performance over an interpolated stimulus space combining both tasks in the same sessions (**Fig. 6d**), ruling out differences in strategies between the two tasks and arguing that rats used the same decision-making system while detecting and categorizing odors.

We found that odor categorization performance is more susceptible to category boundary fluctuations than odor identification (**Fig. 8**). This is due to high stimulus input that always exists in this task, and thus the multiplication in **Equation 4** implies amplification in weight update when sensory evidence is large. Our model is in agreement with previous observations for the categorization task (Uchida and Mainen, 2003; Zariwala *et al.*, 2013). In particular, the much smaller tradeoff between accuracy and reaction times observed in this task is not due to a lack of quality of stimulus input. In fact, our model indicates that signal-to-noise ratio is extremely high for the categorization task (**Fig. 8d**). This suggests that high performance can be achieved with short integration time even for mixtures near 50-50%, particularly compared to the identification task, in which signal-to-noise ratio is highly reduced. Additional Bayes- and RL-DDM simulations do in fact show that performance remains relatively unaltered in mixture categorization with an increase of integration threshold, contrasting with what would be predicted for odor identification (**Fig. S18**). This agrees with the observation that one sniff is enough for maximum performance in mixture categorization (Uchida and Mainen, 2003). Weight fluctuations, which impair performance in a trial-by-trial basis, cannot be filtered out within the integration process. On the other hand, the identification task is highly driven by stimulus noise, which is reflected within the diffusion process, and thus favored by integration in order to make better decisions. We thus conclude that the observation of different speed-accuracy tradeoffs is due to different computational requirements in the two tasks.

We have demonstrated that the simple scenario of detecting a noisy stimulus is insufficient to capture all the details occurring in a two-forced choice task. Two other effects have to be incorporated in order to explain the differences observed here. First, the effect of reward bias, as previously described (Busse *et al.*, 2011; Scott *et al.*, 2015) and second, a novel result of ongoing stimulus-dependent learning (**Fig. 6**). These results show that, given the right conditions, learning can be detrimental for the rats’ performance while categorizing stimuli. Nevertheless, these trial-by-trial dependencies might not be observed in all tasks. In the case of visual discrimination of random dots coherence (Roitman and Shadlen, 2002; Palmer, Huk and Shadlen, 2005) or auditory discrimination of clicks (Brunton, Botvinick and Brody, 2013), for example, the stimuli are lateralized in accordance with the correct decision direction and the decision boundary is set at a natural neutral point, the midline. We hypothesize that these tasks will show reduced updating effects, as the midline boundary represents a strong prior that the subjects have experienced extensively, contrasting with the 50/50 odor mixture separation boundary which is an arbitrary mapping to left/right responses that must be learned.

We have shown ongoing learning compromises the rats’ ability to categorize odors. We have ensured that ongoing learning is not due to incomplete learning as the rats present stable performance over the analyzed data (**Suppl. Figs. 19, 20**). This suggests some other interfering factor or some deliberate strategy of the rats. One possible interpretation is that the rats’ performance is limited by their inability to remember over a very long trial history. An ideal decision maker would learn to set the perfect decision boundary by averaging over all the trials that it has been exposed to. Imperfect memory would imply that the most recent trials will have a larger effect in deciding what to do with a given stimuli (conditional on ongoing learning occurring, *a Ψ* 0, i.e. that the most recent trial is still affecting performance). An alternative interpretation is that the memory is not imperfect but that the animal is constantly learning. The learning could be due to changes in the response of the sensory epithelium or uncontrolled trial to trial variations in the stimulus or experimental trig. Alternatively, the animal might learn because it wrongly assumes that the rules of the task change over time, even though this is not the case. The psychophysics-like experimental paradigm is indeed highly artificial in the sense that outcomes and states are crystallized. It is unlikely that this would be the case in a more naturalistic environment, where, due to environmental dynamics, odors could signal different outcomes, rewards and states over time. A normal, ever-changing environment would imply adaptability and never-ending learning as the optimal strategy. This strategy becomes suboptimal in a static environment, but this may be a small price to pay compare to the cost of stopping learning erroneously when the world is actually dynamic. These results are consistent with a recent proposal that suboptimal inference, as opposed to internal noise, is a major source of behavioral variability (Beck *et al.*, 2012). In this case, the apparent suboptimal inference is the result of assuming that the world is dynamic when, in fact, it is static.

## EXPERIMENTAL PROCEDURES

### Animal subjects

Four Long Evans rats (200-250 g at the start of training) were trained and tested in accordance with European Union Directive 86/609/EEC and approved by Direcção-Geral de Veterinária (DGV) of Portugal. Rats were pair-housed and maintained on a normal 12 hr light/dark cycle and tested during the daylight period. Rats were allowed free access to food but were water-restricted. Water was available during the behavioral session and for 20 minutes after the session at a random time as well as on non-training days. Water availability was adjusted to ensure animals maintained no less than 85% of *ad libitum* weight at any time.

### Testing apparatus and odor stimuli

The behavioral apparatus for the task was designed by Z.F.M. in collaboration with M. Recchia (Island Motion Corporation, Tappan, NY). The behavioral control system (BControl) was developed by Z.F.M, C. Brody (Princeton University) in collaboration with A. Zador (Cold Spring Harbor Laboratory). The behavioral setup consisted of a box (27 × 36 cm) with a panel containing three conical ports (2.5 cm diameter, 1 cm depth). Each port was equipped with an infrared photodiode/phototransistor pair that registered a digital signal when the rat’s snout was introduced into the port (“nose poke”), allowing us to determine the position of the animal during the task with high temporal precision. Odors were delivered from the center port and water from the left and right ports. Odor delivery was controlled by a custom made olfactometer designed by Z.F.M in collaboration with M. Recchia (Island Motion Corporation, Tappan, NY). During training and testing the rats alternated between two different boxes. The test odors were S-(+) and R-(-) stereoisomers of 2-octanol, chosen because they have identical vapor pressures and similar intensities. In the odor identification task, difficulty was manipulated by using different concentrations of pure odors, ranging from 10^-4^ to 10^-1^ (v/v).

The different concentrations were produced by serial liquid dilution using an odorless carrier, propylene glycol (1,2-propanediol). In the odor mixture categorization task, we used binary mixtures of these two odorants at different ratios, with the sum held constant: 0/100, 20/80, 32/68, 44/56 and their complements (100/0, etc.). Difficulty was determined by the distance of the mixtures to the category boundary (50/50), denoted as “mixture contrast” (e.g., 80/20 and 20/80 stimuli correspond to 60% mixture contrast). Choices were rewarded at the left choice port for odorant A (identification task) or for mixtures A/B > 50/50 (categorization task) and at the right choice port for odorant B (identification task) or for mixtures A/B < 50/50 (categorization task). In both tasks, the set of eight stimuli were randomly interleaved within the session. During testing, the probability of each stimulus being selected was the same. For the experiment in **Figs. 2, 4, 5** and **6**, only mixtures with a total odor concentration of 10^-1^ were used. For the experiment in **Fig. 2**, we used the same mixture contrasts with total concentrations ranging from 10^-1^ to 10^-4^ prepared using the diluted odorants used for the identification task. In each session, four different mixture pairs were pseudo-randomly selected from the total set of 32 stimuli (8 contrasts at 4 different total concentrations). Thus, for this task, a full data set comprised 4 individual sessions.

For all the different experiments, four of the eight stimuli presented in each session were rewarded on the left (odorant A, for detection; A/B > 50/50, for categorization) and the other four were rewarded on the right (odorant B, for detection; A/B < 50/50, for categorization). Each stimulus was presented with equal probability and corresponded to a different filter in the manifold.

For the experiments in **Fig. S2** we used two different sets of mixture ratios: 0/100, 17/83, 33.5/66.5, 50/50 in one experiment and 0/100, 39/61, 47.5/52.5, 49.5/50.5 in the second experiment. In the experiment using 50/50 mixture ratios we used two filters both with the mixture 50/50, one corresponding to the left-rewarded stimulus and the other one to the right-rewarded stimulus. Thus, for the 50/50 mixtures, rats were rewarded randomly, with equal probability for both sides.

### Reaction time paradigm

The timing of task events is illustrated in **Fig. 1**. Rats initiated a trial by entering the central odor-sampling port, which triggered the delivery of an odor with delay (*d_odor_*) drawn from a uniform distribution with a range of [0.3, 0.6] s. The odor was available for up to 1 s after odor onset. Rats could exit from the odor port at any time after odor valve opening and make a movement to either of the two reward ports. Trials in which the rat left the odor sampling port before odor valve opening (~4% of trials) or before a minimum odor sampling time of 100 ms had elapsed (~1% of trials) were considered invalid. Odor delivery was terminated as soon as the rat exited the odor port. Reaction time (the Odor sampling duration) was calculated as the difference between odor valve actuation until odor port exit (**Fig. 1**) minus the delay from valve opening to odor reaching the nose. This delay was measured with a photo ionization detector (mini-PID, Aurora Scientific, Inc) and had a value of 53 ms.

Reward was available for correct choices for up to 4 s after the rat left the odor sampling port. Trials in which the rat failed to respond to one of the two choice ports within the reward availability period (~1% of trials) were also considered invalid. For correct trials, water was delivered from gravity-fed reservoirs regulated by solenoid valves after the rat entered the choice port, with a delay (*d_water_*) drawn from a uniform distribution with a range of [0.1, 0.3] s. Reward was available for correct choices for up to 4 s after the rat left the odor sampling port. Trials in which the rat failed to respond to one of the two choice ports within the reward availability period (0.5% of trials) were also considered invalid. Reward amount (*w_rew_*), determined by valve opening duration, was set to 0.024 ml and calibrated regularly. A new trial was initiated when the rat entered odor port, as long as a minimum interval (*d_inter-trial_*), of 4 s from water delivery, had elapsed. Error choices resulted in water omission and a “time-out” penalty of 4 s added to *d_inter-trial_*. Behavioral accuracy was defined as the number of correct choices over the total number of correct and incorrect choices. Invalid trials (in total 5.8 ± 0.8% of trials, mean ± SEM, n = 4 rats) were not included in the calculation of performance accuracy or reaction times (odor sampling duration or movement time).

### Training and testing

Rats were trained and tested on three different tasks: (1) a two-alternative choice odor identification task; (2) a two-alternative choice odor mixture categorization task (Uchida and Mainen, 2003); and (3) a two-alternative choice “odor mixture identification” task. The same rats performed all three tasks and all other task variables were held constant.

The training sequence consisted of: (I) handling (2 sessions); (II) water port training (1 session); (III) odor port training, in which a nose poke at the odor sampling port was required before water was available at the choice port. The required center poke duration was increased from 0 to 300 ms (4 – 8 sessions); (IV) introduction of test odors at a concentration of 10^-1^, rewarded at left and right choice ports according to the identity of the odor presented (1 – 5 sessions); (V) introduction of increasingly lower concentrations (more difficult stimuli) (5 – 10 sessions); (VI) training on odor identification task (10 – 20 sessions); (VII) testing on odor identification task (14 – 16 sessions); (VIII) training on mixture categorization task (10 – 20 sessions); (IX) testing on mixture categorization task (14 – 15 sessions); (X) testing on mixture identification task (12 – 27 sessions) (**Fig. S1**).

During training, in phases V-VII, we used adaptive algorithms to adjust the difficulty and to minimize bias of the animals. We computed an online estimate of bias:

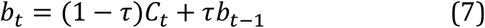

where *b_t_* is the estimated bias in the current trial, *b_t-1_* is the estimated bias in the previous trial, *Ct* is the choice of the current trial (0 if right, 1 if left) and *τ* is the decay rate (*τ* = 0.05 in our experiments). The probability of being presented with a right-side rewarded odor *p* was adjusted to counteract the measured bias using:

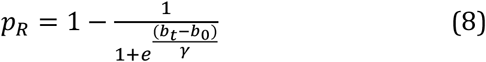

where *b_0_* is the target bias (set to 0.5), and *γ* (set to 0.25) describes the degree of non-linearity. Analogously, the probability of a given stimulus difficulty was dependent on the performance of the animal, i.e., the relative probability of difficult stimuli was set to increase with performance. Performance was calculated in an analogous way as (1) at the current trial but *c_t_* became *r_t_* – the outcome of the current trial (0 if error, 1 if correct). A difficulty parameter, *δ*, was adjusted as a function of the performance,

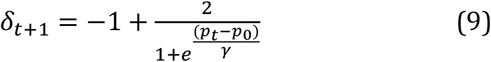

where *p_0_* is the target performance (set to 0.95). The probability of each stimulus difficulty, *φ*, was drawn from a geometric cumulative distribution function (GEOCDF, Matlab)

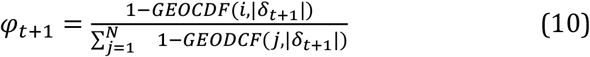

where *N* is the number of stimulus difficulties in the session, and takes a value from 2 to 4 (when *N*=1, i.e. only one stimulus difficulty, this algorithm is not needed); *i* corresponds to the stimulus difficulty and is an integer from 1 to 4 (when *δ* > 0, the value 1 corresponds to the easiest stimuli and 4 to the most difficult one, and vice-versa when *δ* < 0). In this way, when |*δ*| is close to 0, corresponding to an average performance close to 0.95, the distribution of stimuli was close to uniform (i.e. all difficulties are equally likely to be presented). When performance is greater, then the relative probability of difficult trials increased; conversely, when the performance is lower, the relative probability of difficult trials decreased. Training phases VI and VIII were interrupted for both tasks when number of stimulus difficulties *N*=4 and difficulty parameter δ stabilized on a session-by-session basis.

Each rat performed one session of 90-120 minutes per day (250–400 trials), 5 days per week for a period of ~120 weeks. During testing, the adapting algorithms were turned off and each task was tested independently. The data set was collected only after performance was stable (**Fig. S19**) during periods in which the animals showed stable accuracy and left/right bias on both tasks (**Figs. S19, S20**). Throughout the test period, there was variability in accuracy and bias across sessions, but there was no correlation between these performance metrics and session number (accuracy: Spearman’s rank correlation *ρ*=-0.066, P=0.61 for identification, *ρ*=0.16, P=0.24 for categorization; bias: *ρ*=0.104, P=0.27 for both tasks, identification: *ρ*=0.093, P=0.48, categorization: *ρ*=0.123, P=0.37).

### Session bias and choice bias

We fit all psychometric curves by considering the cumulative Gaussian distribution and a lapse rate *l_r_* and minimizing least the least square difference through fminsearch (Matlab). In this way we were able to describe the psychometric curve with two additional parameters, slope (variance) and threshold (mean)(Stevens, 1975; Busse *et al.*, 2011):

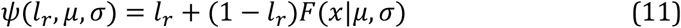

where *F*(*x*|*μ*, *σ*) is a cumulative Gaussian. We defined choice bias as the difference between the values of *ψ* and chance (0.5) at exactly the middle of the stimulus space (*x=0* for the identification task, *x=50%* for the categorization task). We quantified this for sessions (**Fig. S20**) and all trials (**Fig. 6c-d**).

We defined the threshold of such psychometric curve as the indifference point (*I = μ*), which indicates the stimulus difficulty at which the rat has an equal chance of choosing left or right (**Fig. 6a-b**, solid black line). In order to look at the effect of a reward and its interaction with stimulus difficulty on choice bias, we conditioned our analyzed psychometric function to trials that followed a correct choice with a given stimulus *X_T-1_* (in which *T-1* represents the preceding trial).

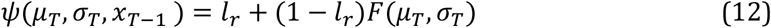

With this analysis we described conditional psychometric curves that differ depending on the outcome and difficulty of the average preceding trial (**Fig. 6a-b**).

Considering the original curve, we quantified the change in choice bias after a given stimulus as the displacement at the indifference point between the two curves (**Fig. 6c-d, Figs. S7 and S8**):

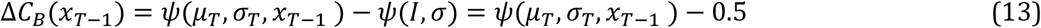

For trials following an error we calculated this change as the displacement at the indifference point between the current curve and two trials back (**Figs. S9** and **S10**). This was done in order to avoid capturing bouts of incorrect trials that might contaminate this analysis.

### Model

#### Drift-diffusion model for decision-making

For a given stimulus with concentrations *c_A_* and *c_B_* we define the accumulated evidence at time *t, e*(*t*). The diffusion process has the following properties: at time t=0 the accumulated combined evidence is zero, *e*(0) = 0; and the momentary evidence *m_i_* is a random variable that is independent at each time step. We discretize time in steps of 0.1ms and run numerical simulations of multiple runs/trials. For each new time step *t*=*nΔt* we generate a new momentary evidence draw:

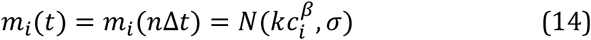

that is, through a normally distributed random generator with a mean of 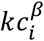, in which we define *k* as the sensitive parameter, and *β* as the exponent parameter. *σ* defines the amount of noise in the generation of momentary evidences. We set *σ* to 1, making 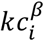 equivalent to the signal to noise ratio for a particular stimuli and respective combination of concentrations (*c_A_ c_B_*). Integrated evidences (*s_1_, s_2_*) are simply the integration of the momentary evidences over time

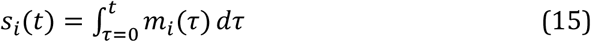

We translate this in our discretized version as a cumulative sum at all time steps, effectively being:

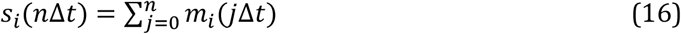

We then define the decision variable accumulated evidence as:

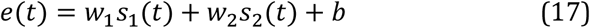

or in it’s discretized version:

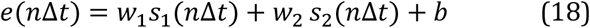

where *w_1_* and *w_2_* are model-dependent combination weights on the accumulated evidence, and *bis* an a-priori decision bias (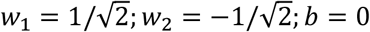 for optimal decisions; 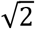 scaling ensures 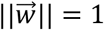). Together, these parameters define slope and offset of the category boundary, which determines the mapping between accumulated evidence and associated choices. We also define the (accumulation) decision bound *θ*(*t*) and make it in most models collapsing over time through either a linear or an exponential decay. Thus, at timestep *nΔt* the bound is either

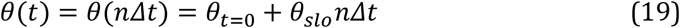

where we define *θ_t=0_* as the bound height at the starting point of integration *t = 0* and *θ_slo_* ≤ 0 as its slope, or

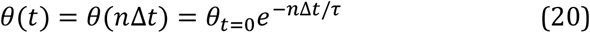

where *τ* ≥ 0 is the bound height’s mean lifetime. The collapse parameters *θ_slo_* and *τ* define the level of urgency in a decision, the smaller it becomes, the more urgent a given decision will become, given rise to more errors (Churchland *et al.*, 2011; Drugowitsch *et al.*, 2012). For models with non-collapsing boundaries we used *θ(t)* = *θ_t=0_*, independent of time. For models with collapsing boundaries, they collapsed linearly, except for RL-DDM, where they collapse exponentially.

Decisions are triggered once the accumulated evidence, *e*(*t*), crosses one of the two decision boundaries {*θ*(*t*),–*θ*(*t*)}. To simulate these decisions, we first simulated a one-dimensional diffusion model that directly uses *e*(*t*) as the diffusing “particle”, and from this reconstructed the higher-dimensional accumulated momentary evidences 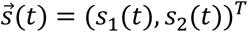. For the onedimensional simulation we used a momentary Gaussian evidence with drift 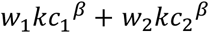 and diffusion variance 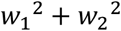 (both per unit time step), corresponding to the moments of *e*(*t*) ‒ *b*. We reintroduce the bias *b* by shifting the boundaries to {*θ*(*t*) – *b*, –*θ(t)* – *b*}. For non-collapsing boundaries we simulated accumulation boundary crossings using a recently developed, fast, and unbiased method (Drugowitsch, 2016). For collapsing boundaries, we simulated these boundary crossing by Euler integration in *Δt* = 0.001*s* time-steps, and set the final *e(t)* to lie on the crossed boundary to avoid overshooting that might arise due to time discretization. In both cases, we defined the decision time *t_d_* as the time when crossing occurred, and the choice in trial *k* by

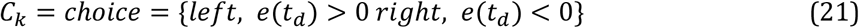

To recover the higher-dimensional accumulated momentary evidences at decision-time, 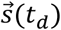, we sampled those from the two-dimensional Gaussian 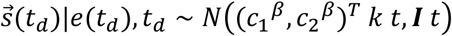 (i.e., unbounded diffusion), subject to the linear decision boundary constraint 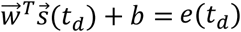, using the method described in Simpson, Turner and Pettitt, 2008.

In order to capture <100% accuracy in easy trials and systematic and consistent choice biases, we introduced an additional “lapse” component with lapse rate *l_r_* and lapse bias *l_b_* to the model. The lapse rate *l_r_* determined the probability with which the choice is not determined by the diffusion model, but is instead drawn from a Bernoulli distribution that chooses “right” with probability *l_b_* and “left” with probability 1 — *l_b_*. Model fits revealed small lapse rates, and lapse biases close to 0.05 (**Suppl. Table 1** and **2**). These lapse rates are typically needed for this type of models and have been hypothesized in the past to be due to effects of attention and/or exploration (Wichmann and Hill, 2001).

Lastly, the reaction time for a particular trial was simulated by adding a normally distributed non-decision time variable with mean *t_ND_* and standard deviation 0.1 *t_d_* to the decision-time arising from the diffusion model simulations (Palmer, Huk and Shadlen, 2005),

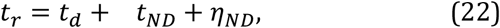

where 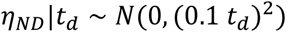 models the stochasticity of the non-decision time. Without weight and bias learning (that is, when fixing 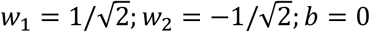), the base model with a non-collapsing has the following six parameters: sensitivity (*k*), exponent (*β*), nondecision time mean (*t_ND_*), initial bound height (*θ_t=0_*), lapse rate (*l_r_*), and lapse bias (*l_b_*). A collapsing bound introduces one additional parameter, which is the boundary slope (*θ_slo_*) for linearly collapsing boundaries, or the boundary mean lifetime (*τ*) for exponentially collapsing boundaries.

#### Drift-diffusion model with Bayesian reward bias and stimulus learning – Bayes-DDM

The following provides an overview of the Bayesian model that learns stimulus combination weights, reward biases, or both. A complete description of the model and its derivation can be found in (Drugowitsch and Pouget, 2018). We first focus on weight learning, and then describe how to apply the same principles to bias learning. The model assumes that there are true, latent combination weights 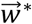 that the decision-maker can’t directly observe, but aims to infer based on feedback on the correctness of his/her choices. To ensure continual learning, these latent weights are assumed to slowly change across consecutive trials *k* and *k* + 1 according to a first-order autoregressive process,

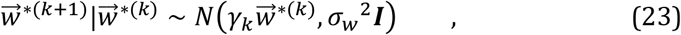

with weight “leak” 0 ≤ *γ_k_* < 1, ensuring that weights remain bounded, and weight diffusion variance 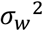, ensuring a continual, stochastic weight change. This process has zero steady-state mean and a steady-state variance of 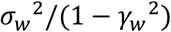 for each of the true weight components, which we used as the decision-maker’s prior 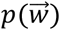 over the inferred weight vector 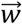.

For each sequence of trials that we simulated, the decision-maker starts with this prior in the first trial and updates its belief about the weight vector in each subsequent trial in two steps. We will describe these two steps on hand of making a choice in trial *k,* receiving feedback about this choice, updating one’s belief, and then moving on to the next trial *k + 1.* Before the first step in trial *k*, the decision-maker holds the “prior” belief *p*(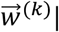 past information) = 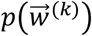 that is implicitly conditional on all feedback received in previous trials 1,…, *k* – 1. The decisionmaker then observes some sensory evidence, accumulates this evidence, commits to choice *C_k_* with decision time *t_k_* and accumulated momentary evidences 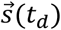. After this, the correct choice *C_k_^*^* ∈ {–1,1} (-1 for “left”, 1 for “right) is revealed, which, in our 2-AFC setup is the same as telling the decision-maker if choice *C_k_* was correct or incorrect. The Bayes-optimal way to update one’s belief about the true weights upon receiving this feedback is given by Bayes’ rule,

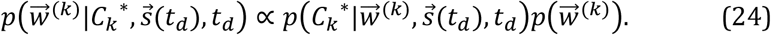

Unfortunately, the functional form of the likelihood 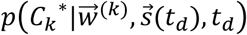 does not permit efficient sequential updating of this belief, but we have shown elsewhere (Drugowitsch and Pouget, 2018) that we can approximate the above without considerable performance loss by assuming that the posterior (and, by induction, also the prior) is Gaussian. Using prior parameters 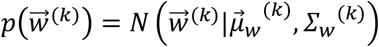 and posterior parameters 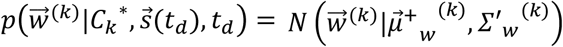 yields the update equations

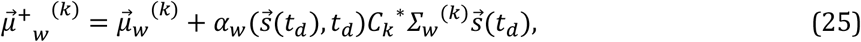

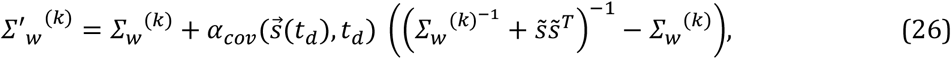

with learning rates

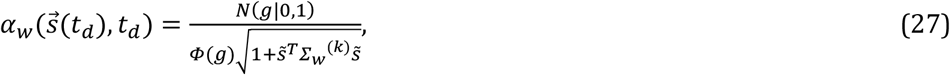

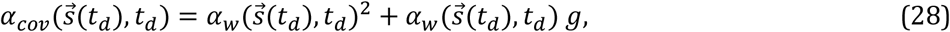

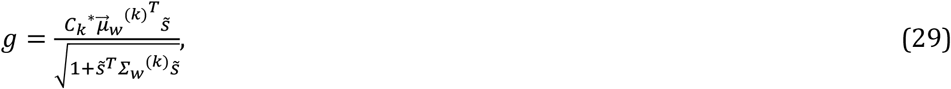

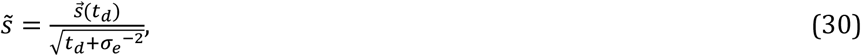

where *Φ*(·) is the cumulative function of a standard Gaussian, and where *σ_e_^2^* is a variance that describes the distribution of decision difficulties (e.g., odor intensities) across trials, and which we assume to be known by the decision-maker. In the above, *g* turns out to be a quantity that is closely related to the decision confidence in trial *k.* Furthermore, both learning rates, *α_w_* and *α_cov_* are strongly modulated by this confidence, as follows: they are small for high-confidence correct decisions, moderate for low-confidence decisions irrespective of correctness, and high for high-confidence incorrect choices. A detailed derivation, together with more exploration of how learning depends on confidence is provided in Drugowitsch and Pouget, 2018.

Once the posterior parameters have been computed, the second step follows. This step takes into account that the true weights change across consecutive trials, and is Bayes-optimally captured by the following parameter updates:

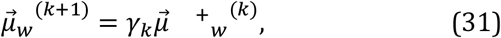

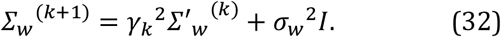

These parameters are then used in trial *k* + 1. Overall, the Bayesian weight learning model has two adjustable parameters (in addition to those of the base decision-making model): the assumed weight leak (*γ_w_*) and weight diffusion variance (*σ_w_^2^*) across consecutive trials.

Let us now consider how similar principles apply to learning the bias term. For this we again assume a true underlying bias *b*\* that changes slowly across consecutive trials according to

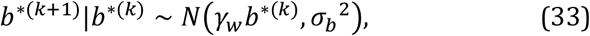

where the leak *γ_w_* is the same as for 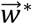, but the diffusion *σ_b_^2^* differs. As we show in Drugowitsch and Pouget, 2018, the bias can be interpreted as a per-trial a-priori bias on the correctness on either choice, which brings it into the realm of probabilistic inference. More specifically, this bias can be implemented by extending the, until now two-dimensional, accumulated momentary evidences 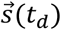 in each trial, by an additional, constant element. An analogous extension of 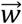 adds the bias term to them, until now two-dimensional, weight vector. Then, we can perform the same Bayesian updating of the, now three-dimensional, weight vector parameters as described weights, to learn weights and the bias simultaneously. The only care we need to take is to ensure that, in the second step, the covariance matrix elements associated with the bias are updated with diffusion variance *σ_b_^2^* rather than *σ_w_^2^*. Overall, a Bayesian model that learns both weights and biases has three adjustable parameters: the assumed weight and bias leak (*γ_w_*), the weight diffusion variance (*σ_w_^2^*), and the bias diffusion variance (*σ_b_^2^*). A Bayesian model that only learns the bias has two adjustable parameters: the assumed bias leak (*γ_w_*), and the bias diffusion variance (*σ_b_^2^*).

#### Drift-diffusion model with heuristic reward bias and stimulus learning – RL-DDM

Rather than using the Bayesian weight and bias update equations in their full complexity, we also designed a model that captures their spirit, but not their details. This model does not update a whole distribution over possible weights and biases, but instead only works with point estimates, which take values 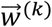 and *b*^(*k*)^ in trial *k*. After feedback *C_k_*^*^ ∈ {–1,1} (as before, -1 for “left”, 1 for “right”), the model updates the weight according to

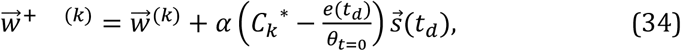

where *α* is the learning rate. Note that, for rapid decisions (i.e., *t_d_ ≈ 0*), we have |*e*(*t_d_*)| ≈ *θ_t=0_*, such that the residual term in brackets is zero for correct choices, such that learning only occurs for incorrect choices. For slower choices and collapsing boundaries, we will have |*e*(*t_d_*)| < *θ_t=0_*, such that the residual will be non-zero even for correct choices, promoting weight updates for both correct and incorrect choices. Considering that decision confidence in the Bayesian model is generally lower for slower choices, this learning rule again promotes learning rates weighted by confidence: fast, high-confidence choices result in no weight updates for correct choices, and large weight updates for incorrect choices, whereas low, low-confidence choices promote moderate updates irrespective of the correctness of the choice, just as for the Bayes-optimal updates. To ensure a constant weight magnitude, the weights are subsequently normalized by

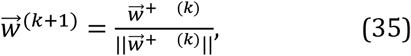

to form the weights for trial *k* + 1.

Bias learning takes a similar flavor, using the update equation

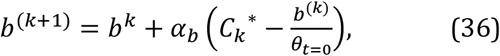

where *α_b_* is the bias learning rate. In contrast to weight learning, this update equation does not feature any confidence modulation, but was nonetheless sufficient to capture the qualitative features of the data. Overall, this learning model added two adjustable parameters to the base decision-making model: the weight learning rate (*α*), and the bias learning rate (*α_b_*).

#### Alternative learning heuristics

To further investigate if a confidence-modulated learning rate was required, we designed models that did not feature such confidence weighting. For weight learning, they used the delta rule

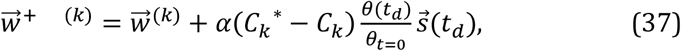

where *α* is the learning rate, and whose weight updated is, as before, followed by the normalization 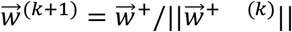. Here, we assume the same encoding of make choice *C_k_* and correct choice *C_k_^*^*, that is, *C_k_* ∈ {–1,1} (-1 for “left”, 1 for “right”), such that the residual in brackets is only non-zero if the choice was incorrect. In that case, the learning rate is modulated by boundary height, but no learning occurs after correct choices.

The bias is learned similarly, using

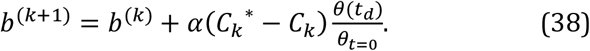

Overall, this results in one adjustable parameter in addition to the base decision-making model: the learning rate (*α*).

#### Drift-diffusion model with reward bias and stimulus weight fluctuations

To test if random weight and bias fluctuations are sufficient to capture the across-task differences, we also fit a model that featured such fluctuations without attempting to learn these weights from feedback. Specifically, we assumed that, in each trial, weights and biases where drawn from

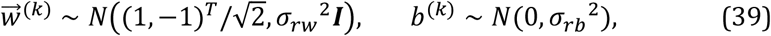

which are normal distributions centered on the optimal weights and bias values, but with (co)variances *σ_rw_^2^I* and *σ_rb_^2^*. We adjusted these (co)variances to best match the data, leading to two adjustable parameters in addition to those of the base decision model: the weight fluctuation variance (*σ_rw_^2^*), and the bias fluctuation variance (*σ_rb_^2^*).

#### Model fitting

We found the best-fitting parameters for each model by log-likelihood maximization (Palmer, Huk and Shadlen, 2005). Due to collapsing bounds and (for some models) sequential updates of weights and biases, we could not directly use previous approaches that rely on closed-form analytical expressions (Ratcliff, 1978) for fitting diffusion models with non-collapsing boundaries. Instead, for any combination of parameters, we simulated the model responses to a sequence of 100,000 trials with stimulus sequence statistics matching those of the rodent experiments for the conditions that we were interested in fitting. These responses were used to compute summary statistics describing model behavior, which were subsequently used to evaluate the log-likelihood of these parameters. We computed the log-likelihood in two ways, first by ignoring sequential choice dependencies, and second by taking such dependencies into account. All model simulations were performed as described further above. We did not explicitly simulate the stochasticity of the non-decision time, but instead included this stochasticity as an additional noise-term in the likelihood function (not explicitly shown below). To describe how we computed the likelihood of model parameters *φ* without taking sequential dependencies into account, let index *m* denote the different task conditions (i.e., a set of odor concentrations for odors A and B), and let *n_m_* be the numbers of observed trials for this condition in the rodent data that we are modeling. For each condition *m*, we approximate the response time distributions by Gaussians, using *t_m_* and *σ^2^_t,m_* to denote the observed mean response time and variance (across trials). Furthermore, let *P_c,m_* be the observed probability of making a correct choice in that condition. The corresponding model predictions for parameters *φ,* extracted from model simulations, are denoted 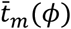 and 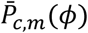. With this, we computed the likelihood of responses times by

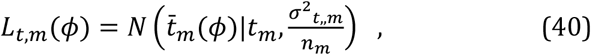

which is the probability of drawing the predicted mean reaction time from a Gaussian centered on the observed mean and with a variance that corresponds to the standard error of that mean. The likelihood of the choice probabilities was for each condition computed by

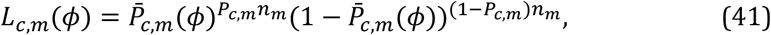

which is the probability of drawing the observed number of correct and incorrect choices with the choice probabilities predicted by the model. The overall log-likelihood is found by summing over the per-condition log-likelihoods, resulting in

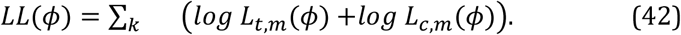

To evaluate the log-likelihood that takes into account sequential choice dependencies, we computed the reaction time likelihoods, *L_t,m_*(*θ*), as before, but changed the choice probability likelihood computation as follows. For trials following correct choices, we computed the choice probability likelihood separately for each stimulus combination given the previous *and* the current trial, thus taking into account that psychometric curves depend on the stimulus condition of the previous trial (**Fig. 6a-b**). Due to the low number of incorrect trials for certain conditions, we didn’t perform this conditioning on the previous trial’s condition when computing the choice probability likelihoods after incorrect choices, but instead computed the likelihood across all trials simultaneously.

For both ways of computing the log-likelihood, we found the parameters that maximize this log-likelihood by use of the Subplex algorithm as implemented in the NLopt library (Steven G. Johnson, The NLopt nonlinear-optimization package, http://ab-initio.mit.edu/nlopt). In some cases we performed the fits without taking into account the sequential choice dependencies, and then predicted these sequential choice dependencies from the model fits (e.g., **Fig. 6c-d**). In other cases (e.g., for some model comparisons), we performed the model fits while taking into account sequential dependencies. The specifics of the model fits are clarified in the main text. The best model fits and respective parameters can be found in **Suppl. Tables 1 and 2**.

#### Model comparison

For comparison between different models with different number of parameters we use Bayesian information criterion (BIC) for model selection (Schwarz, 1978). For each model we calculate the BIC (Wit, Heuvel and Romeijn, 2012):

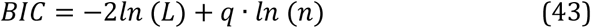

where *q* is the number of free parameters fitted by the model and *n* the number of trials that we fitted. Each model has a BIC associated to it. We compared different models by first converting the BIC score into a log10-based marginal likelihood, using –0.5*B1C/ln(10)*, and then compared models by computing the log10-Bayes factor as the difference between these marginal likelihoods. These differences dictate the explanatory strength of one model in relation to the other. The model with the larger marginal likelihood is preferred and the evidence in favor is decisive if the log-10 difference exceeds 2.

To ensure that our analysis is not driven by the strong parameter number penalty that BIC applies, we performed the same analysis using the Akaike information criterion (AIC) and its corrected version (AICc), but found qualitatively no change in the results. All different model comparisons can be found in **Fig. S4**.

In **Fig. S4** we compared the following models. Models denoted simply “DDM” were diffusion models with optimal weights, 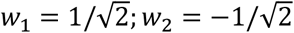. Models denoted “Bayes-DDM” learned their weights as described in the Bayes-DDM section. The “Random weights” models used weights that were stochastically and independently drawn in each trial (see Stimulus weight fluctuations section). The “Delta rule” models learned their weights by the delta rule. The “Full RL-DDM” model used the learning rules described in the RL-DDM section. Only “lapse” variants of these models included the lapse model components. Decision boundaries were constant except for the “collapsing boundary” model variants. The bias was foxed to *b* = 0, except for the “Full RL-DDM” model and “bias” variants. In these bias variants, the biases (but not necessarily the weights, depending on the model) were learned as described in the Bayes-DDM section, except for the “Delta rule” models, for which bias learning was described in the Alternative learning heuristics section. In **Fig. S4**, all models are compared to the Bayes-DDM model that learns both weights and the bias, includes a lapse model, and has collapsing boundaries.

#### Weights fluctuation analysis

As the RL-DDM model reaches a decision it has access to two variables, amount of evidence at the bound and the decision time *t_d_*. For better understanding the dynamics immediately before the multiplication of the weights, we looked at the combination of sensory evidence (*s_1_ s_2_*) for each simulated trial. For each trial *j* there is a noisy sensory evidence trajectory (integration layer from **Fig. 6**). This means that by the end of trial *j* we can compute the mean drift rates that gave to rise to a decision:

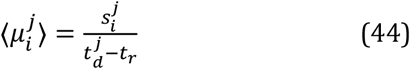

Each group in **Fig. 8a** and **b** has been segregated taking into account the Mahalanobis distance, as each line represents the distance of D=1 for a particular stimulus set.

Considering the integrated evidence of Equation 13 and combined with the choice function of Equation 21 we see that

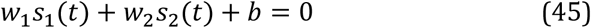

Should represent the separation line between the two stimuli, and thus we can rewrite Equation 45 as:

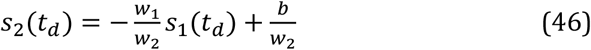

Considering the straight-line equation *y* = *mx* + *i* we see that in our integrated evidence plots the boundary separation can be drawn with slope 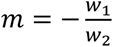 and intercept 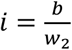.

Stimulus weight fluctuation should then have an impact in the slope of the boundary line separating the classification between left and right stimuli, and *b* should influence the origin intercept on that stimulus representation (**Fig. 8**). Considering the data points simulated for 100.0 trials, we analyzed the effect of slope fluctuation in error rates. That is, how many errors would the model create by having a particular value of m, for both the identification and categorization task (**Fig. 8**).

#### Analysis

All the behavioral and statistical analysis, as well as all fitting, were performed in Matlab®. The different models were implemented and fitted in Julia v0.6.

## ACKNOWLEDGEMENTS

We thank Joseph Paton, Marta Moita, Alfonso Renart, Tim Hanks, Chris Summerfield and the Mainen and Pouget Laboratories for helpful discussions on the work presented here. In particular, we would like to thank Jeff Beck and Ingmar Kanitschider on feedback regarding model conceptualization and implementation.

This work was supported by grants from the Champalimaud Foundation (MIV, AGM, EDW, ZFM), European Research Council (Advanced Investigator Grant 250334, ZFM), Fundação para a Ciência e a Tecnologia (AGM, MIV), Human Frontier Science Program (Grant RGP0027/2010, ZFM & AP), Simons Foundation (Grant 325057, ZFM & AP), and the University of Geneva (AP).

## AUTHOR CONTRIBUTIONS

AGM, MIV and ZFM designed the experiments. AGM, JD, ZFM and AP designed the models. MIV conducted the experiments with assistance from AGM. MIV, AGM, JD and EDW analyzed the data. AGM and JD implemented the models. AGM, JD, MIV, AP and ZFM wrote the manuscript.

