## Supplementary material for "The impact of learning on perceptual decisions and its implication for speed-accuracy tradeoffs"

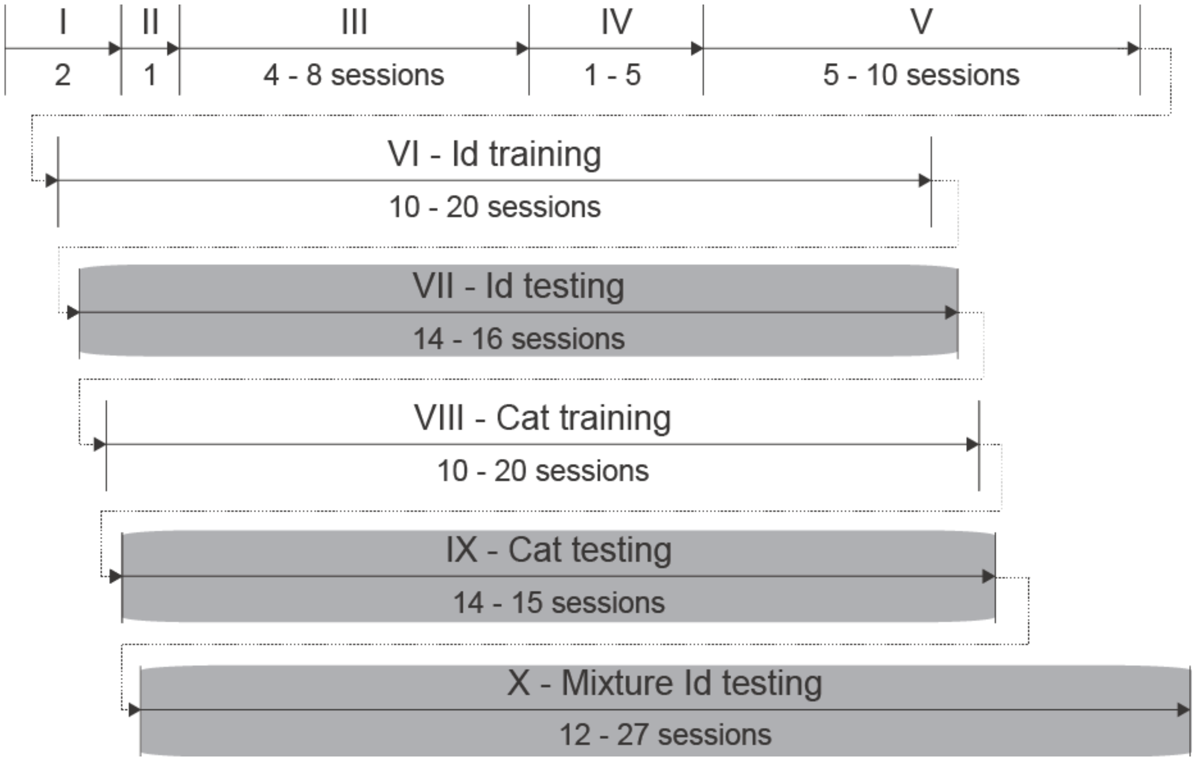


**Figure S1.** Diagram showing the different steps of rats' training and task testing. (I) Handling (2 sessions). (II) Water port training (1 session). (III) Odor port training (4 – 8 sessions). (IV) Pure odor training (1 – 5 sessions). (V) Introduction of lower concentrations (5 – 10 sessions). (VI) Odor identification training (10 – 20 sessions). (VII) Odor identification testing (14 – 16 sessions). (VIII) Mixture categorization training (10 – 20 sessions); (IX) Mixture categorization testing (14 – 15 sessions). (X) Mixture identification testing (12 – 27 sessions). Each session represents a different day. Grey boxes highlight the analyzed data. See Experimental Procedures for more details.

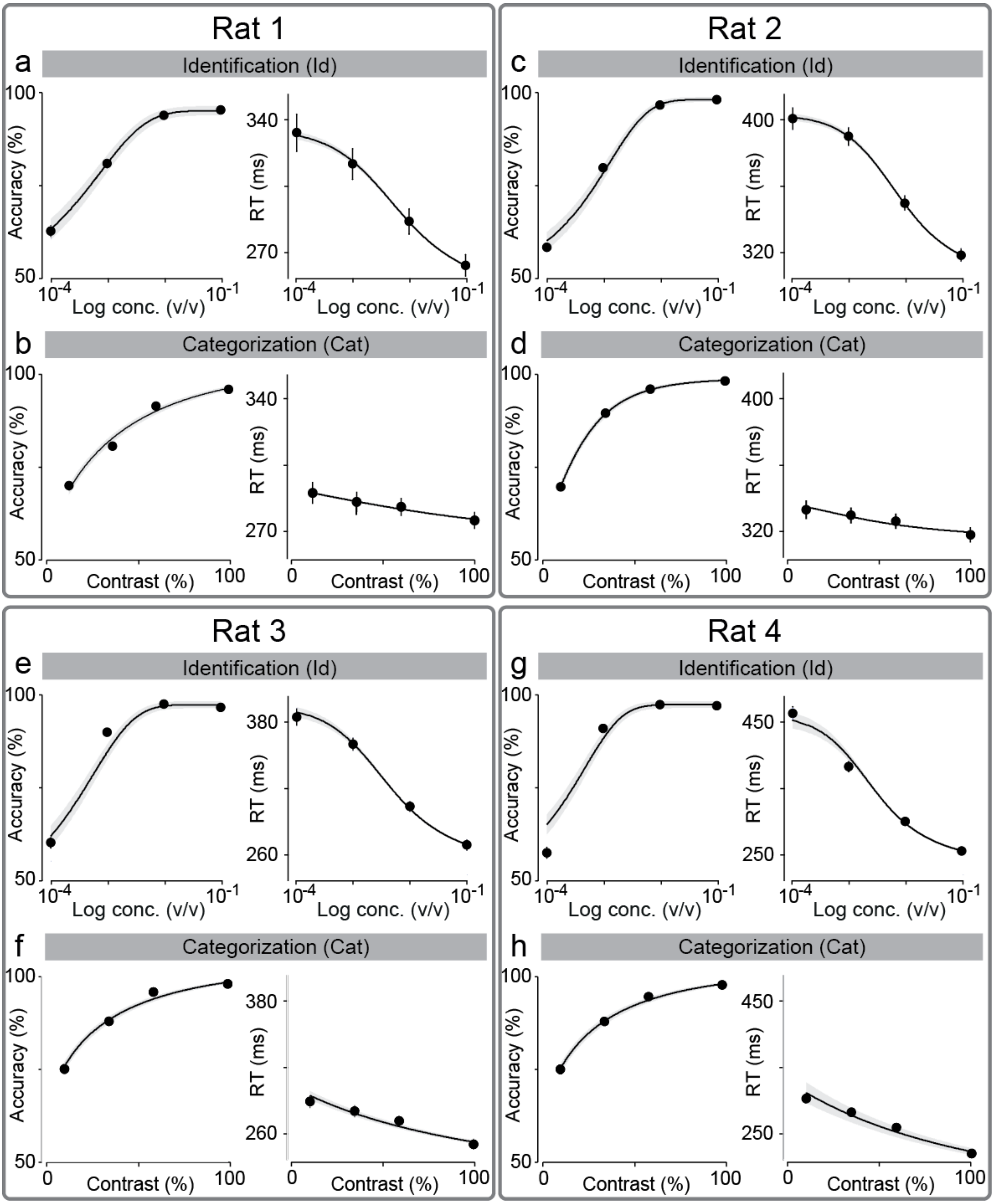


**Figure S2.** Individual behavioral data and DDM fits. **(a,b,e,f)** Behavioral data and fitting results for accuracy and RT in identification task for each individual rat as a function of odor concentration. **(c,d,g,h)** Behavioral data and fitting results for accuracy and RT in categorization task for each individual rat as a function of mixture contrast. Error bars are mean ± SEM over trials. Solid black lines depict each fitted DDM.

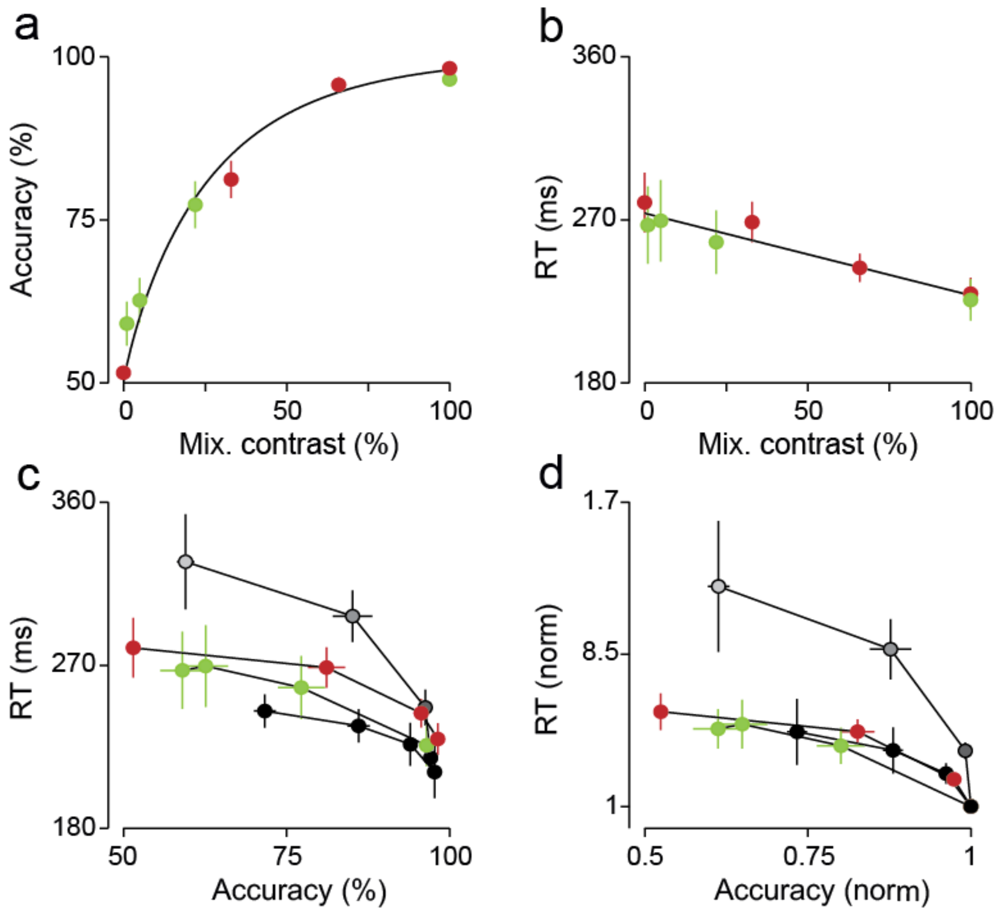


**Figure S3.** Odor mixture categorization with lower contrast stimuli. **(a)** Mean accuracy **(b)** and mean of the median Reaction Times (RTs) plotted as a function of mixture contrast. Red circles represent the set of stimuli: 0, 33, 66, 100% contrast; green circles: 1, 5, 22, 100% contrast. These sets of stimuli were run in different sessions. For the first dataset (red circles), RTs showed an increase of 50 ± 19 ms, from 229 ± 9 to 280 ± 16 ms, from the easiest to the most difficult stimuli ($F(3,12) = 3.98$, $P = 0.04$, one-way ANOVA); for the second dataset (green circles), RTs increased by 41 ± 24 ms, from 226 ± 11 to 267 ± 21 ms ($F(3,12) = 1.17$, $P > 0.3$, one-way ANOVA). **(c)** Mean of median RTs as a function of mean accuracy. Dot shading represents odor concentration, with highest concentration corresponding to the darkest shade; black circles represent the set of stimuli: 12, 36, 60, 100% contrast; red and green circles are the same as in (a, b). **(d)** Normalized mean of median RTs as a function of normalized mean accuracy. Error bars are mean ± SEM (n = 4 rats).

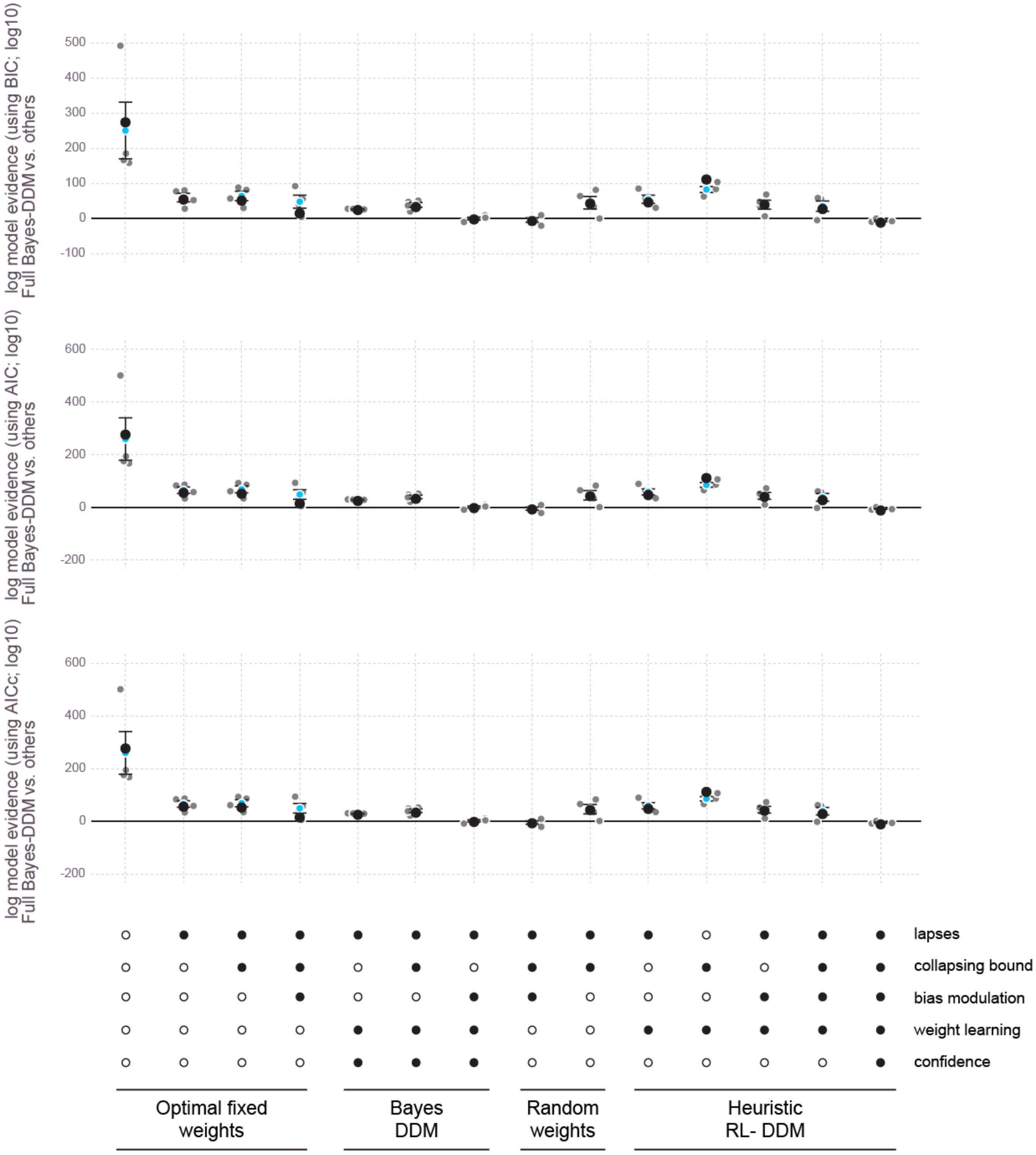


**Figure S4.** Relative fit quality (log difference) for all 14 tested models against the best-fitting model, the full Bayes-DDM. We used Bayes Information Factor (BIC), Akaike information criterion with (AICc) and without (AIC) correction to evaluate fit quality. Model quality is evaluated by considering the objective function fitted on the mean psycho- and chronometric functions (see Experimental Procedures for details). Here we depict 15 different variations of DDM models, with different combinations that encompass the presence or absence of: collapsing bounds, initial bias, lapses, random weights, delta rule updating and confidence. The lower the value of the difference, the better the model fits the mean data. The grey dots show the Bayes factor for individual rats, and the black dots the Bayes factors for the fits using the accumulated data across all rats (adjusted for the increased number of trials). Blue dots show the mean ± SEM seen across rats (n=4). For more details on the relevant models please refer to Experimental Procedures.

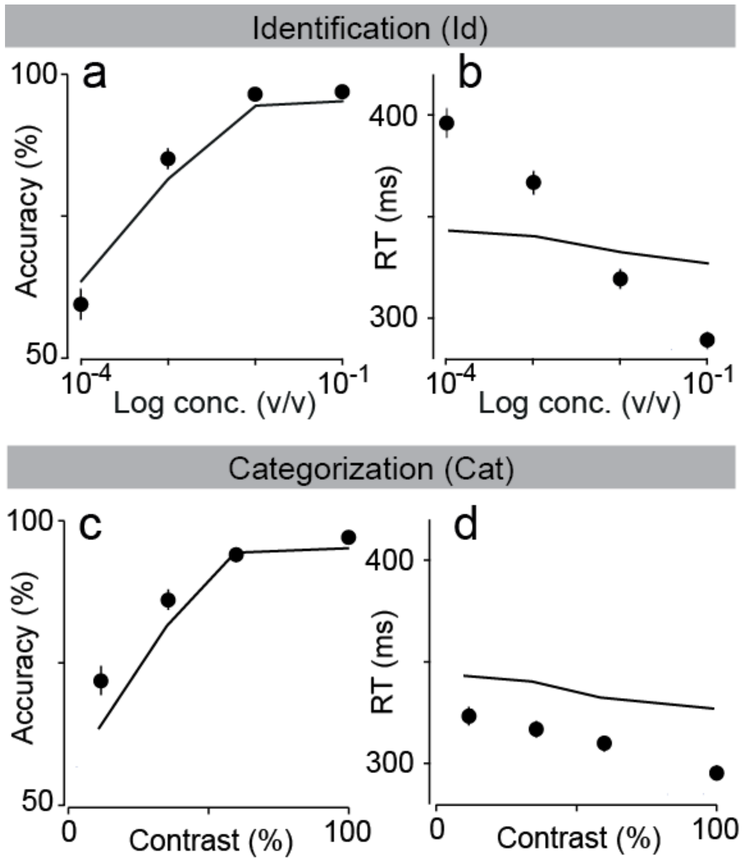


**Figure S5.** Failure to simultaneously fit performance on identification and categorization tasks with unique DDM. In **Fig. 4** the Drift-Diffusion Model was fitted to the identification task data and then used to predict the categorization task behavior (solid lines), and vice-versa (dashed lines). Here we fit both tasks simultaneously (expanded log-likelihood function, with 32 points to fit). Choice accuracy (fraction of correct trials) and reaction times in identification task **(a,b)** and categorization task **(c,d)**. Error bars show SEM trials per rat. Note that these fits assume that six parameters are conserved between the two tasks.

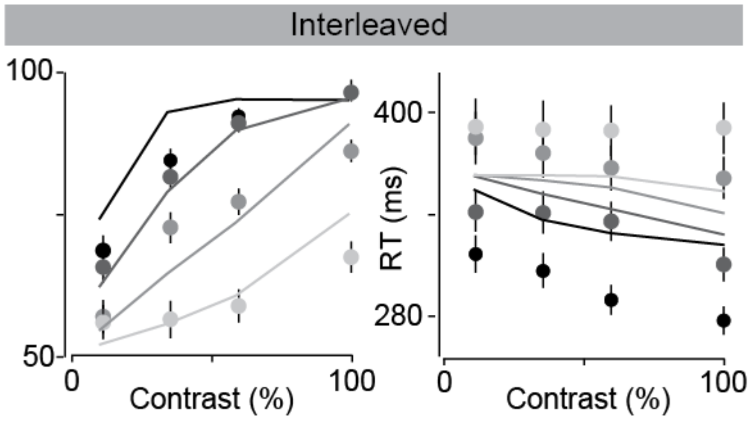


**Figure S6**. Failure to fit interleaved condition data with a collapsing bound DDM. The same DDM model used in **Fig. 4** was used to fit the interleaved condition presented in **Fig. 3**. The model was not able to fit the data adequately, in particular for RTs (right panel). Error bars show SEM across average trials per rat.

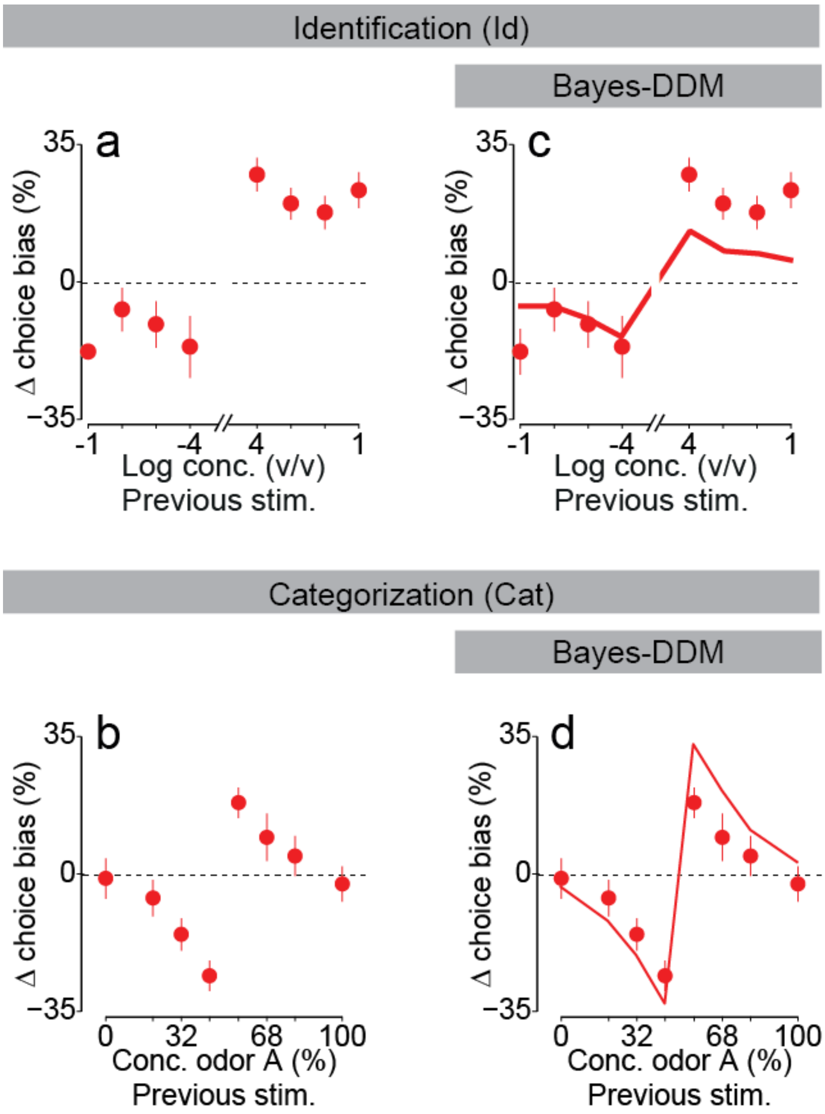


**Figure S7.** Full updating curves for change in choice bias. Change in choice bias after a correct choice and a given stimulus. Data presented for identification **(a,c)** and categorization task **(b,d)**. Note the symmetric change is bias, dependent if the rewarded stimulus was following a left choice (positive bias) or a right choice (negative bias). Regardless, the magnitude of the effect is modulated by the nature of the current stimulus (for identification task), and by the magnitude of the previous trial difficulty (for categorization task). Plotted are also the Bayes-DDM predictions for the updating curves for both tasks (from **Fig. 5**). Error bars show 95% confidence intervals of the measurement.

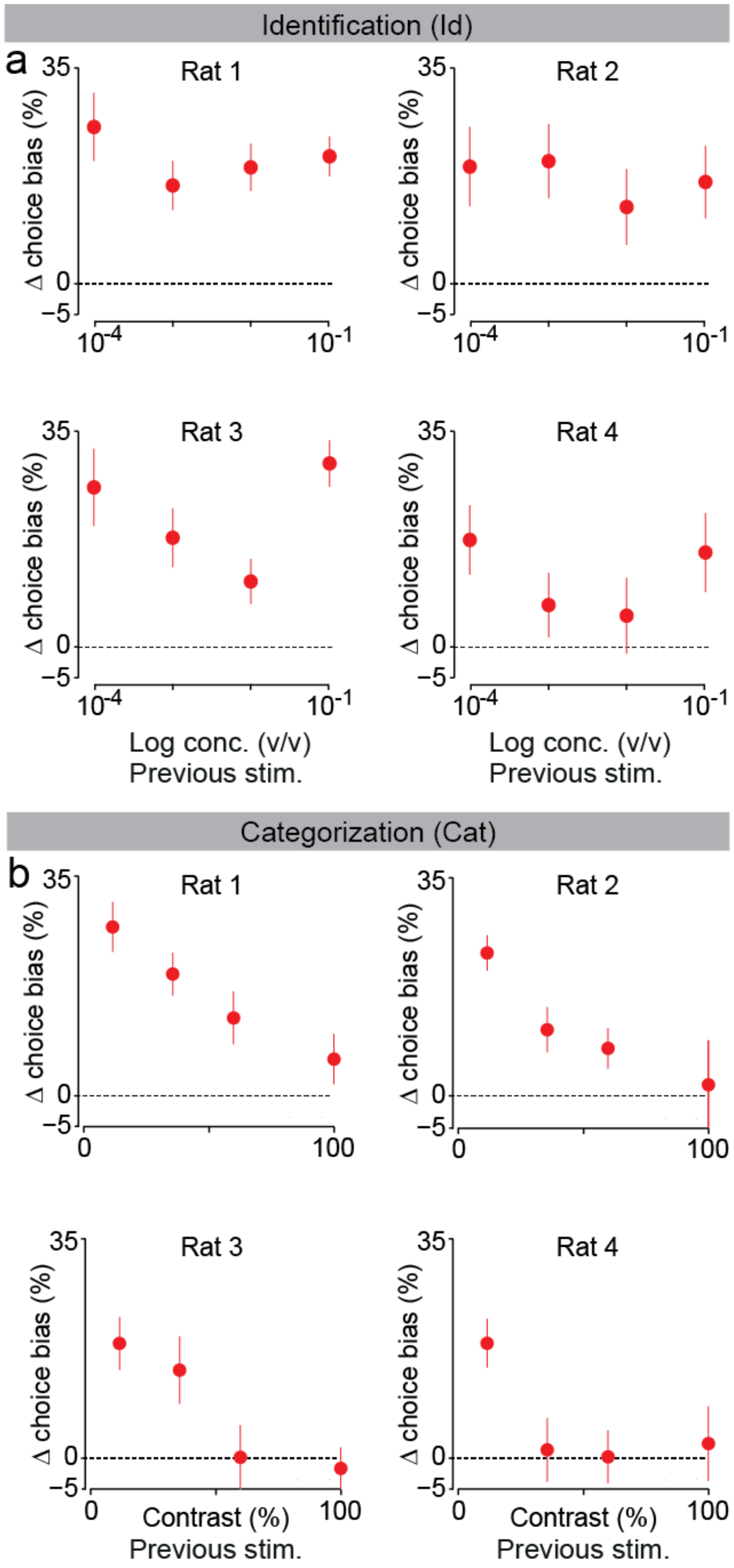


**Figure S8.** Updating curves for change in choice bias after rewarded trials for all individual rats and tasks. Change in choice bias after rewarded choice and a given stimulus. Data presented for identification **(a)** and categorization task **(b)** for all 4 rats. All rats show no effect of stimulus difficulty in change of choice bias for identification task (rat 1, $F(3,59) = 2.5$, $P = 0.069$; rat 2, $F(3,59) = 2.34$, $P = 0.083$; rat 3, $F(3,55) = 2.46$, $P = 0.072$; rat 4, $F(3,51) = 1.54$, $P = 0.214$). For categorization task all rats showed stimulus difficulty modulation in change of choice bias (rat 1, $F(3,55) = 4.39$, $P < 0.01$; rat 2, $F(3,55) = 11.65$, $P <$ 10-5; rat 3, $F(3,55) = 22.73$, $P <$ 10-4;rat 4, $F(3,51) = 3.35$, $P < 0.05$). Error bars show SEM across sessions.

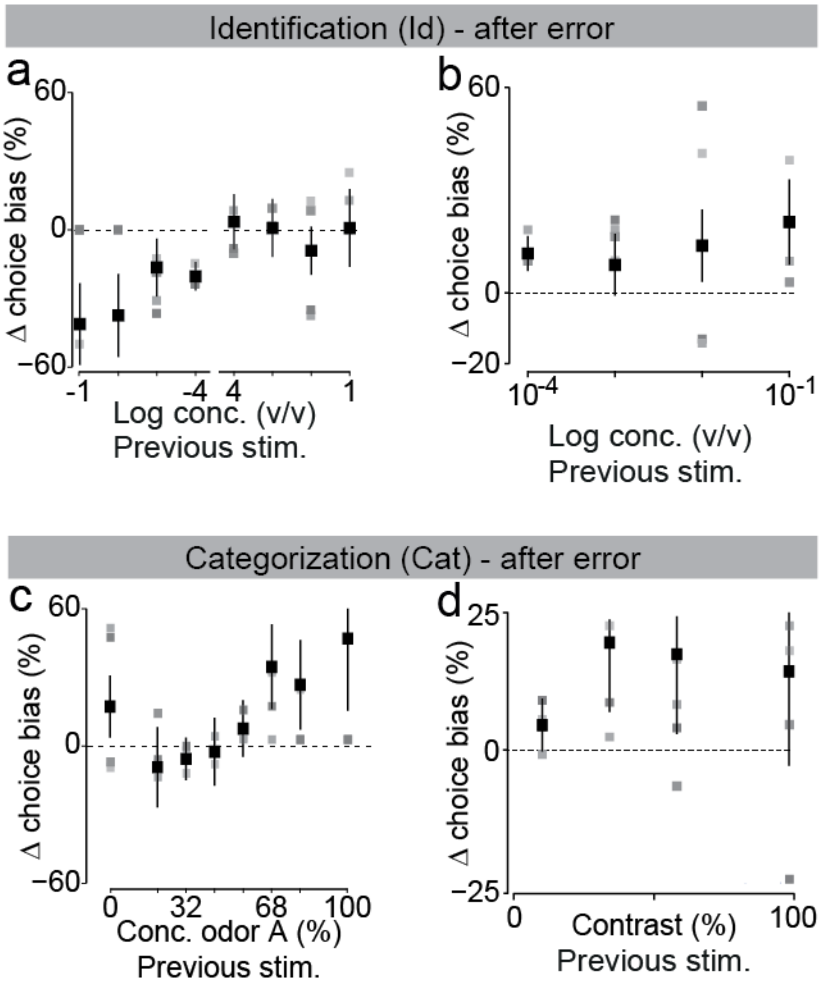


**Figure S9**. Updating curves for change in choice bias after an error response in both tasks. Change in choice bias after an error and a given stimulus. Data presented for identification **(a,b)** and categorization task **(c,d)**. (a,b) Change in choice bias modulated by stimulus concentration. (c,d) Change in choice bias modulated by stimulus difficulty (contrast). Black squares represent mean change when considering all trials from all rats taken together, and the error bars the standard error of the measurement considering a 95% confidence interval. Gray squares show the obtained measurement when considering each rat individually.

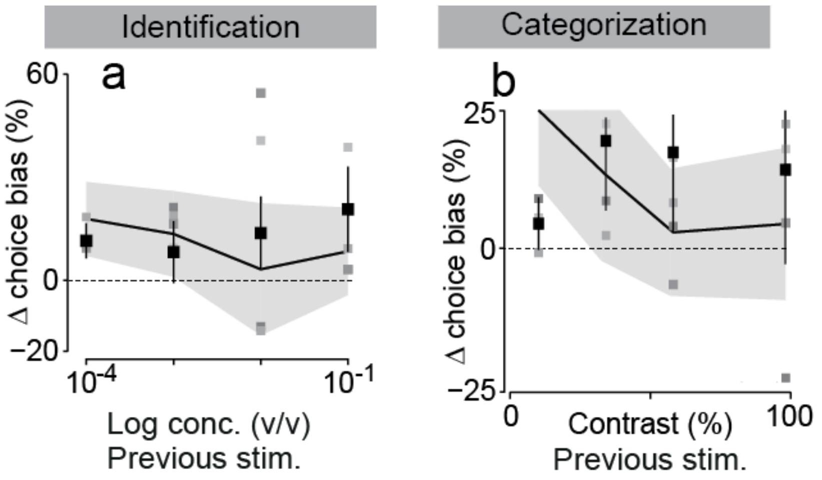


**Figure S10**. Collapsed updating curves for change in choice bias after error response in both tasks comparison. Change in choice bias for identification **(a)** and categorization task **(b)**. Black squares represent mean change when considering all trials from all rats taken together, and the error bars the standard error of the measurement considering a 95% confidence interval. Gray squares show the obtained measurement when considering each rat individually. Black line depicts best fitted Bayes-DDM prediction, and grey area the 95% confidence interval of the predicted effect corrected by the true number of trials seen in the data (not simulated trials). The expected variability from the model makes this analysis troublesome, indicating that, in the model fitting procedure, conditional psychometric curves after incorrect don't constrain the model as well as following a reward. This is probably due to the much larger number of correct trials.

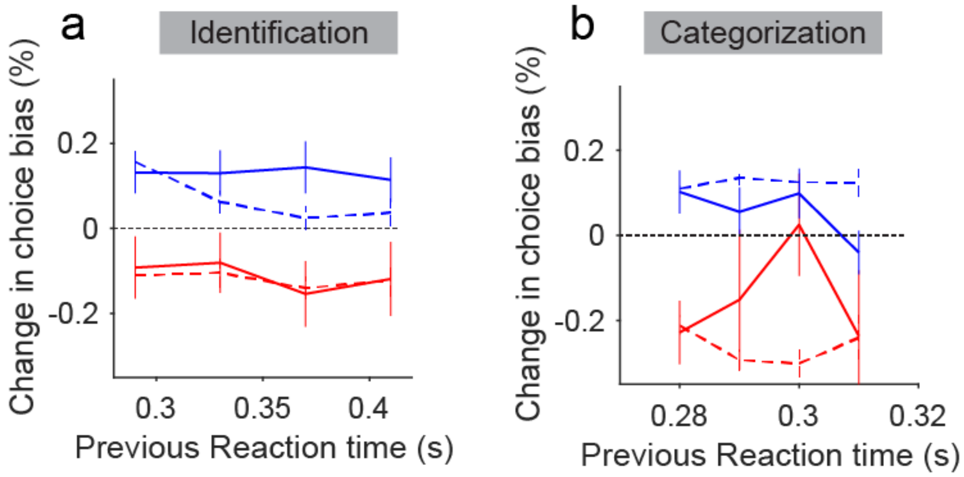


**Figure S11.** Updating curves for change in choice bias after correct and error response in both tasks considering reaction time of previous trial. Change in choice bias for identification **(a)** and categorization task **(b).** Blue lines represent changes after a correct choice, and red after an error choice. Plotted are the observed changes seen in the data (solid lines) and the predicted changes by Bayes-DDM (dashed lines). Each choice bias was calculated considering four time windows for each task. Trials are binned considering the odor sampling duration for that particular trial. Note that even though there is a strong dependence on stimulus difficulty for updating (**Fig. S10**), choice bias change is fairly stable regardless of reaction time. This indicates that, in these tasks, reaction time is not the measurement of confidence, despite the correlation with stimulus difficulty.

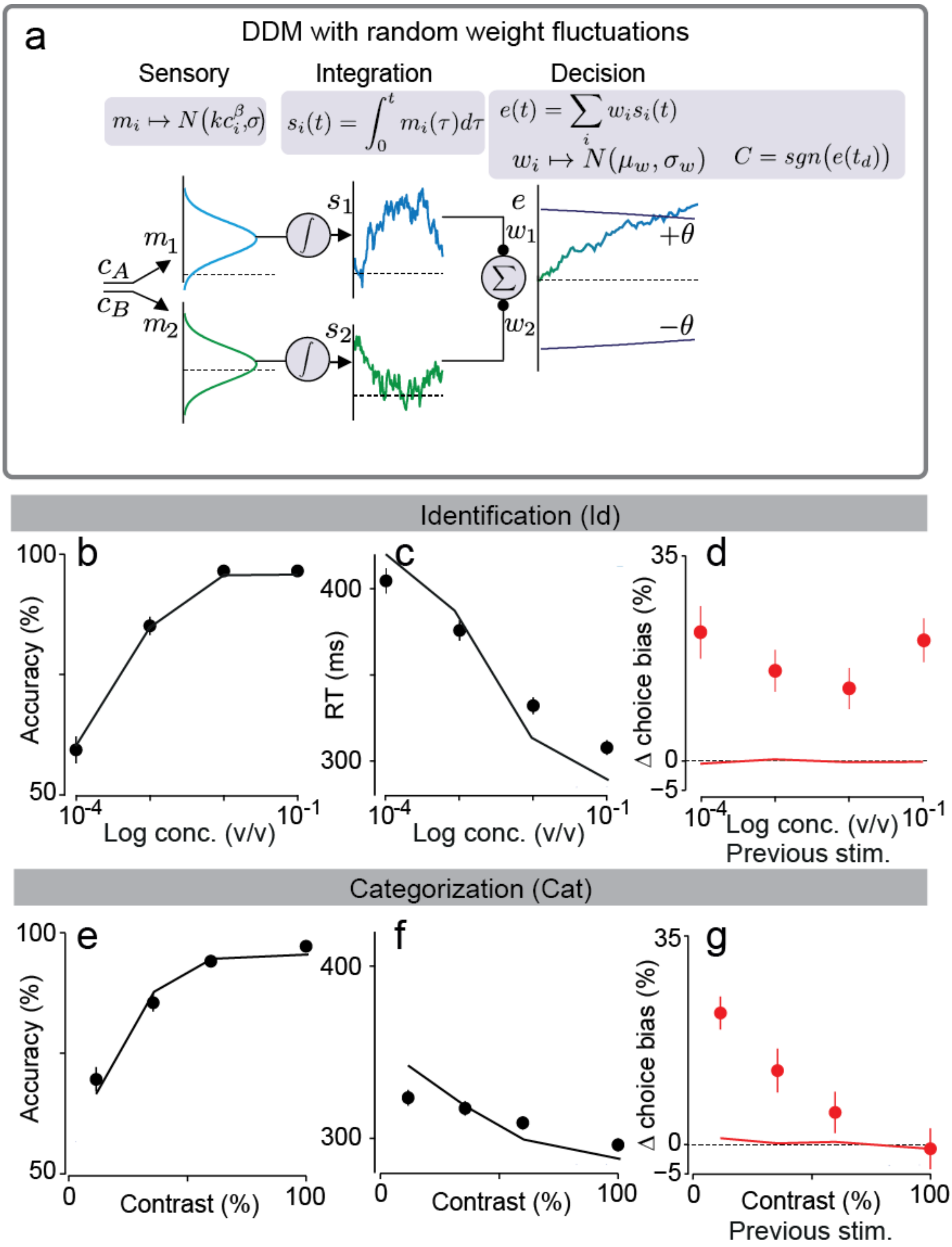


**Figure S12.** Random with random weight fluctuations. **(a)** DDM is expanded to add variability in weight map from stimulus to decision. This is done by varying on a trial-by-trial basis the weights that a transform the stimulus representation into the Decision layer. The magnitude of this effect is set by the standard deviation of the gaussian curve contaminating the weights. This model presents 9 parameters (See Experimental Procedures). Choice accuracy (fraction of correct trials) and reaction times in identification task **(b,c)** and categorization task **(e,f)**. Solid black lines show fits of the model to both tasks. **(d,g)** Change is choice bias after a given choice and rewarded trial for identification (d) and categorization (g) tasks. Note that this model does a good job by fitting both tasks in all psych- and chrono-metric curves. This makes sense as this model effectively adds trial-by-trial variability. However, this model predicts no correlation between trials, as the gaussian noise injected at weight level does not depend on the previous outcome (d,g). Error bars show SEM across rats and trials.

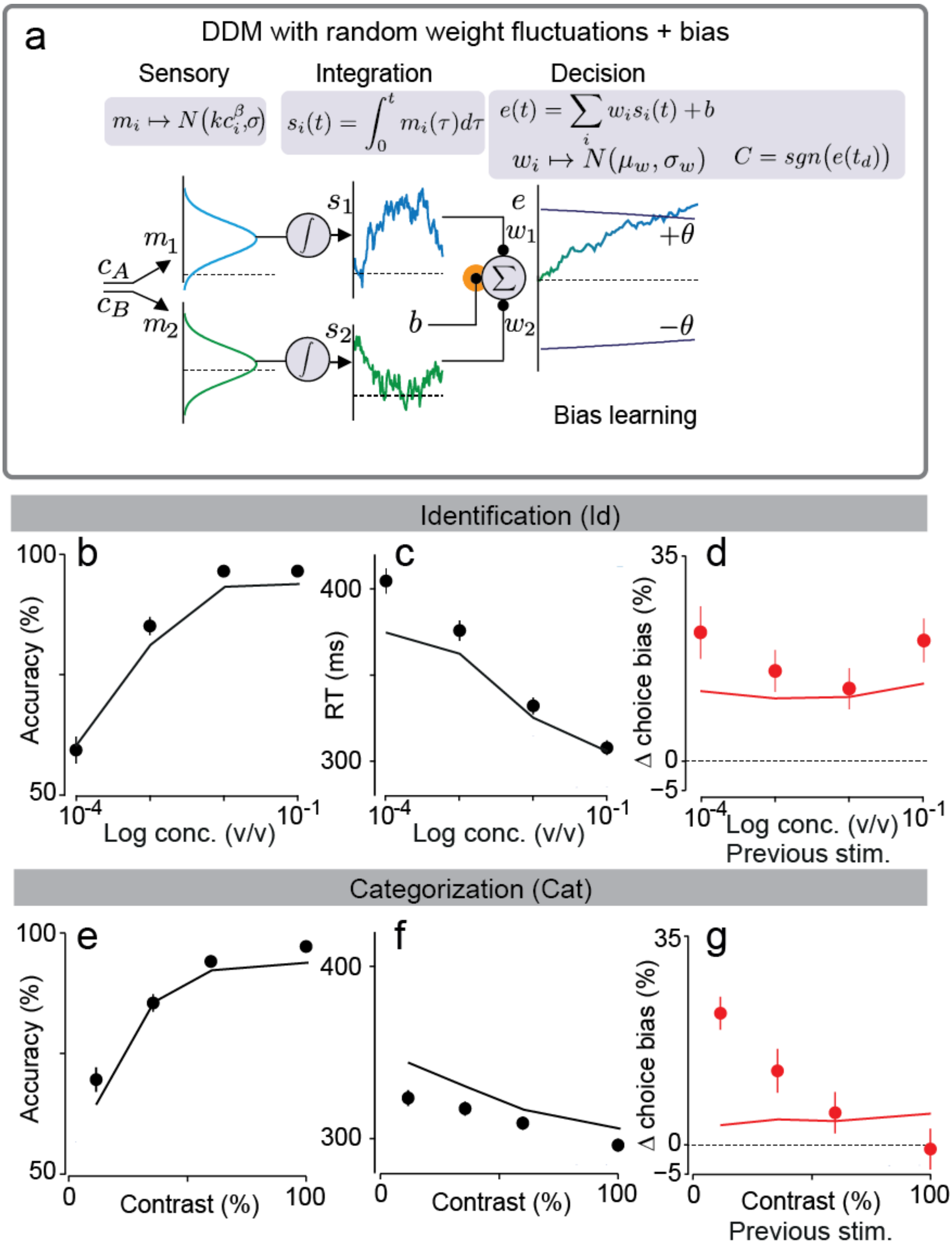


**Figure S13.** Random with random weight fluctuations plus bias. **(a)** Random weights DDM is expanded by adding bias learning, as done in Bayes-DDM. This model presents 10 parameters (See Experimental Procedures). Choice accuracy (fraction of correct trials) and reaction times in identification task **(b,c)** and categorization task **(e,f)**. Solid black lines show fits of the model to both tasks. **(d,g)** Change is choice bias after a given choice and rewarded trial for identification (d) and categorization (g) tasks. Note that this model does a good job by fitting both tasks in all psycho- and chrono-metric curves. This makes sense as this model effectively adds trial-by-trial variability. It also predicts an adequate response of change in choice bias for the identification task (d). However, this model predicts no modulation of difficulty in choice bias for categorization (g). Error bars show SEM across rats and trials.

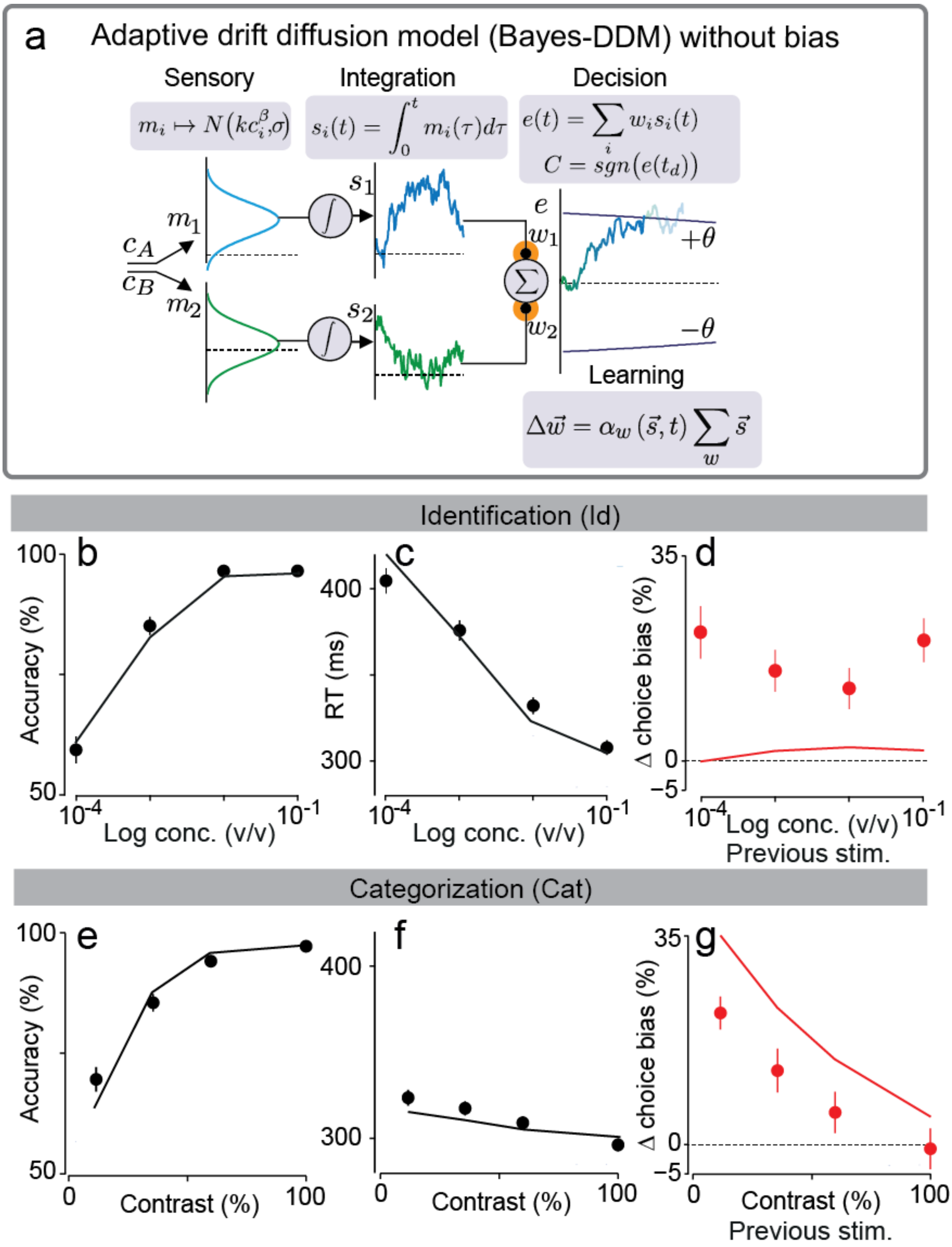


**Figure S14.** Bayes-DDM without bias. **(a)** This model is in all details similar to the Bayes-DDM except for the absence of bias modulation on a trial-by-trial basis (initial starting position). Note that this effectively means that Bayes-DDM's starting position is always at zero evidence. This model presents 9 parameters (See Experimental Procedures). Choice accuracy (fraction of correct trials) and reaction times in identification task **(b,c)** and categorization task **(e,f)**. Solid black lines show fits of the model to both tasks. **(d,g)** Change is choice bias after a given choice and rewarded trial for identification (d) and categorization (g) tasks. Note that this model does a good job by fitting both tasks in all psycho- and chrono-metric curves, as well as choice bias updating for the categorization task (g). However, it lacks a change in choice bias for the identification task (d), thus justifying the presence of a stimulus-independent bias in Bayes-DDM.

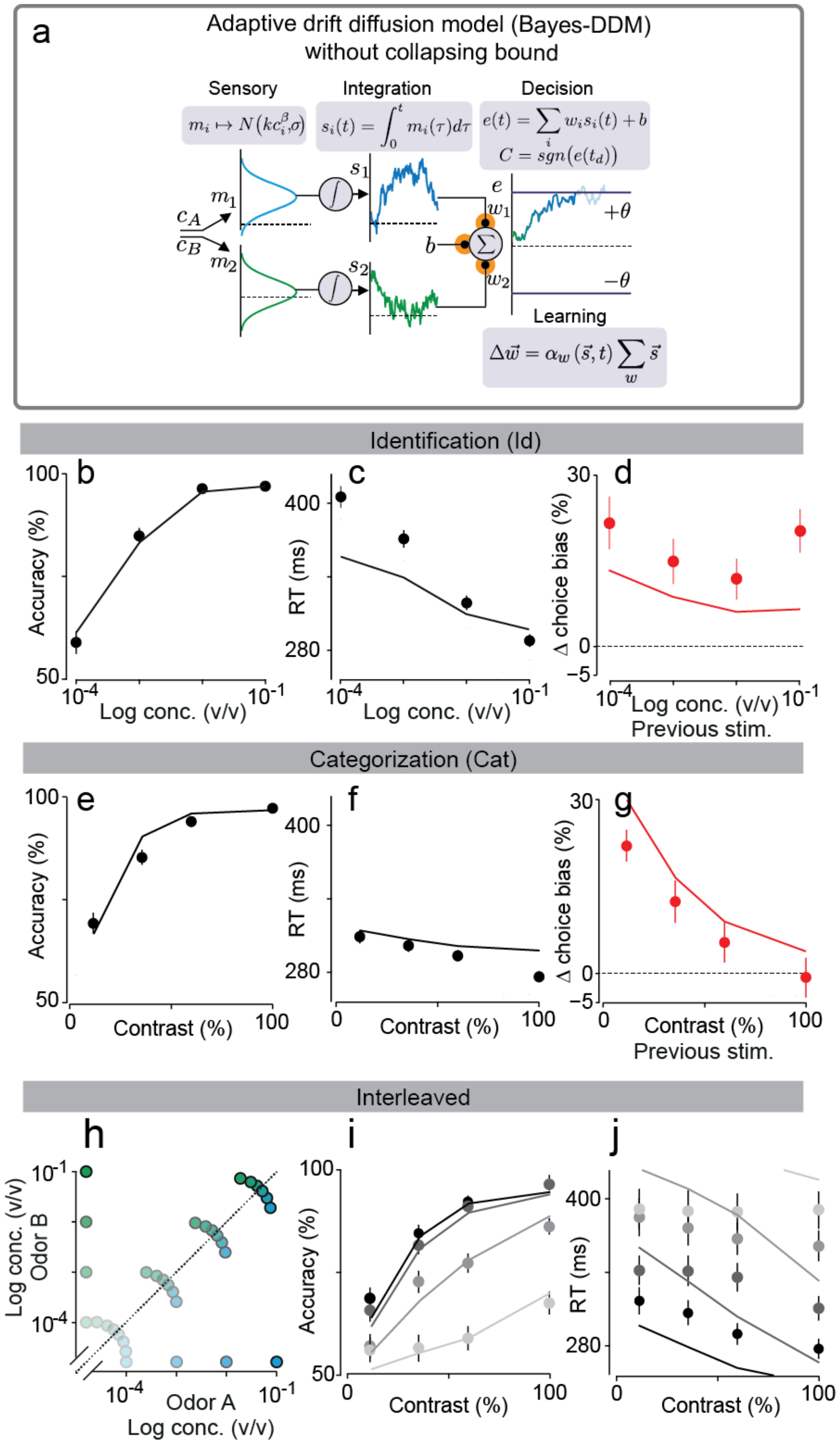


**Figure S15.** Bayes-DDM without collapsing bounds. **(a)** For this model we eliminated the collapsing bound component of Bayes-DDM. THis model presents 9 parameters (See Experimental Procedures). Choice accuracy (fraction of correct trials) and reaction times in identification task **(b,c)** and categorization task **(e,f)**. Solid black lines show fits of the model to both tasks. **(d,g)** Change is choice bias after a given choice and rewarded trial for identification (d) and categorization (g) tasks. **(h,i,j)** Choice accuracy (fraction of correct trials) and reaction times in interleaved task. Solid black lines show fits of the model to this task. This model is one of the best fitting model in our BIC comparison (**Fig. S4**). It does a very accurate job in describing the behavioral data obtained for both taks runned separately (a-g). However, it does present an increased range of reaction times when fitting the behavioral data for the interleaved condition (j). Note that this data was not quantified in the BIC analysis, which explains why this model ranked well in our model comparison.

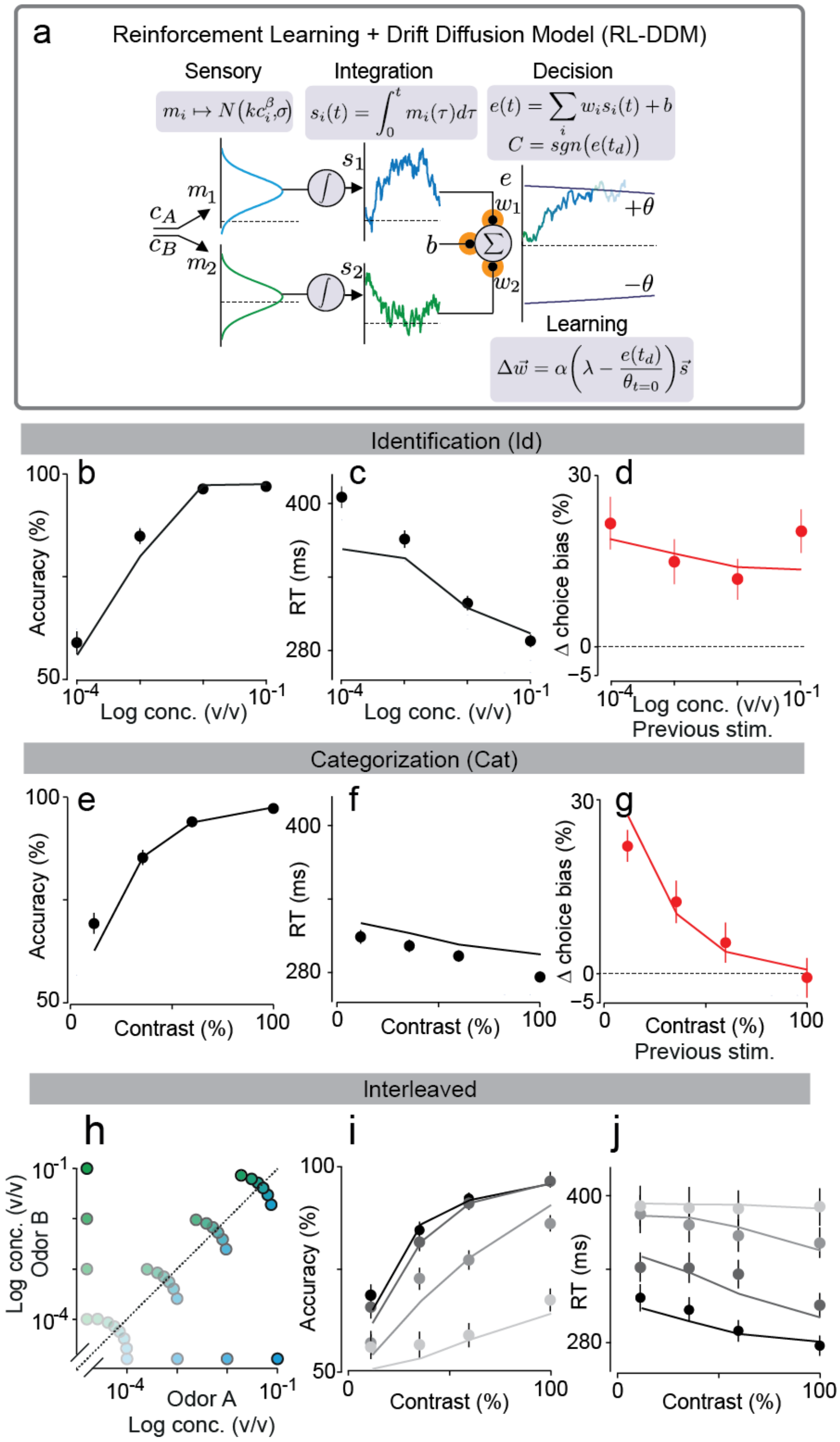


**Figure S16.** Adaptive heuristic DDM with bias and stimulus learning explains identification and categorization task simultaneously. **(a)** Integration DDM is even further expanded with the addition of changing stimulus weights, $w_1$ and $w_2$, and trial-by-trial reward dependent bias *b*. These weights are then combined with the integrated momentary evidences ($s_1$,$s_2$) plus the offset set by the bias b. After each trial the model updates stimulus weights according to the obtained outcome through a delta-learning rule. This model has 9 parameters (see Experimental Procedures). **(b,c,e,f)** Choice accuracy (fraction of correct trials) and odor sampling duration in identification task (b,c) and categorization task (e,f). Solid black line represents model with 10 parameters fitted to identification and categorization task. Error bars are mean ± SEM over trials and rats. **(d,g)** Change in choice bias plotted as a function of the previous stimuli. All four different odor concentrations for identification task (d); and all mixture sets for categorization (g). Red points correspond to behavioral data, and solid red line to the predicted change from the fitted model. Error bars represent ± SEM over 4 rats. (**h,i,j**) Choice accuracy (fraction of correct trials) and reaction times in interleaved task. Solid black lines show fits of the model to this task. This model is one of the best fitting model in our BIC comparison (**Fig. S4**), as well as one of the best predicting the changes in choice bias and fitting the interleaved data.

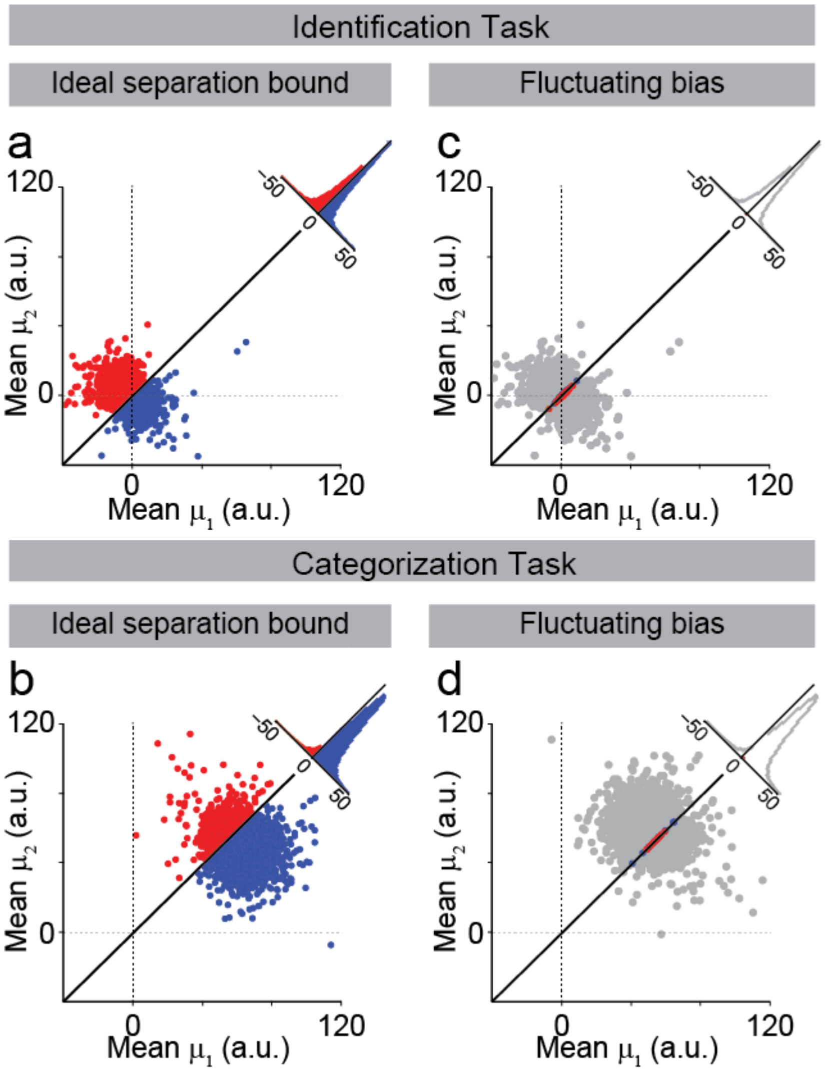


**Figure S17.** Bias fluctuations affect categorization and identification equally. **(a,b)** Mean drift rates for the most difficult left-decision choice in the case of the Bayes-DDM model. In the case of the identification task this is the $10^{-4}$/0 stimuli (a); for the categorization task the stimulus is the 56%/44% mixture (b). Blue signal the correct classified choices and red the incorrect. Considering the ideal separation bound we show the projected histograms for the difference ⟨μ1⟩-⟨μ2⟩. (**c,d**) Same as (a,b) but now with fluctuating bias depicted as the intercept of the category bound i=b. Grey area indicates the area of bias fluctuations that is equivalent to 1 standard deviation of the bias fluctuations for both identification (c) and categorization (d) tasks. Blue indicates trials that were originally incorrect in (a,b) but became correct, and red indicates trials that became incorrect but originally correct. Light grey dots indicate answers that remained unchanged from (a,b). Histograms quantify the population for each of the four populations of dots. Note that the change is minimal, in particular when compared to the fluctuations seen for stimulus weight updating (**Fig. 8**).

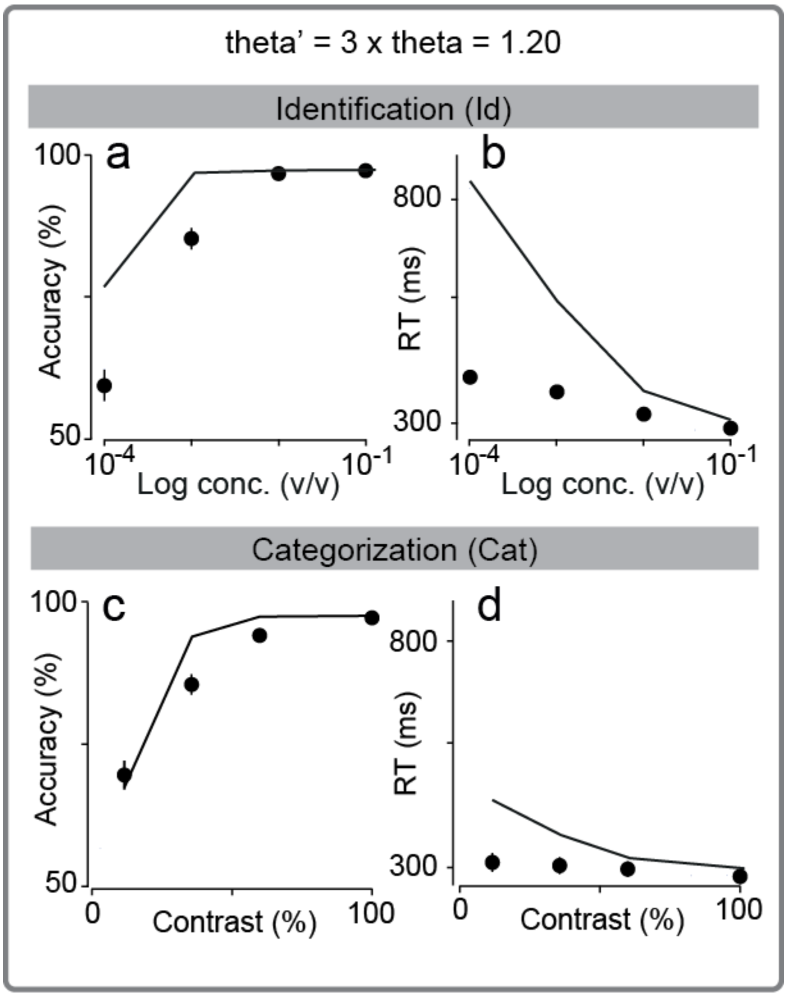


**Figure S18.** Bayes-DDM predictions with diffusion threshold increase for both tasks. **(a,b)** Psycho- and chronometric predictions for identification task considering a three-fold increase in threshold in Bayes-DDM. **(c,d)** Same as (a,b) but for categorization task. Error bars show SEM across rats and trials. Lines show predictions from the Bayes-DDM.

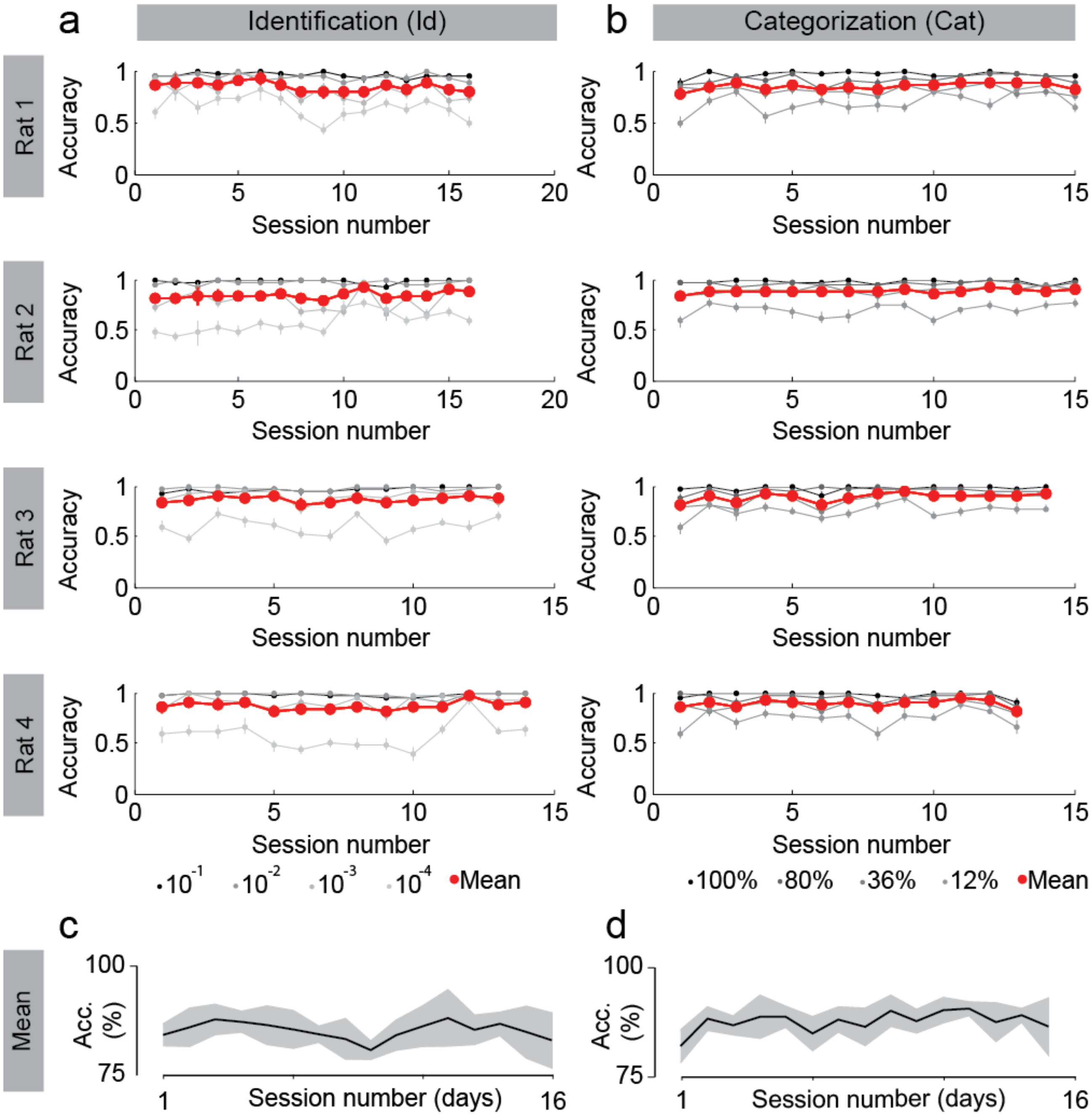


**Figure S19.** Session-by-session stable performance over testing days. Performance for each individual rat plotted on a session-by-session basis. Data is presented for both identification **(a)** and categorization task **(b)**. Depicted are all the sessions used in the analysis. Each difficulty is presented separately, with color codes as depicted on the figure (bottom). Mean accuracy for all trials is shown in red. Overall performance was stable despite local fluctuations for different difficulties, in particular for difficult trials. Error bars show proportion SE for each session and each individual rat, across trials. **(c,d)** Mean session-by-session change in overall accuracy over rats for both identification (c) and categorization (d) tasks. Solid black lines are the mean and shaded grey the SEM over the 4 rats. Each session was conducted on a different day.

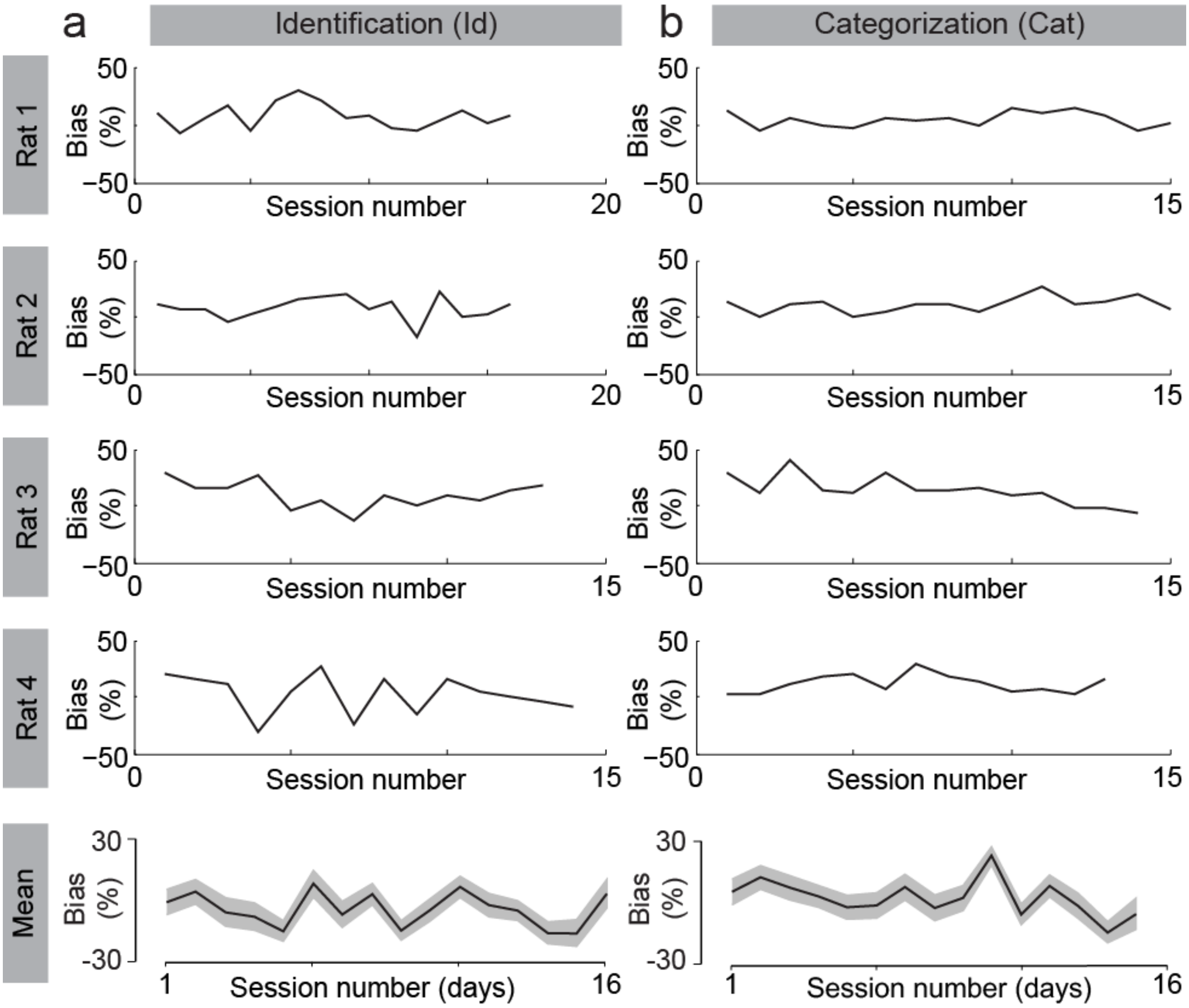


**Figure S20.** Session-by-session performance bias fluctuations over testing days. Session bias (tendency to go left, B>0, or right, B<0) fitted for each individual rat and respective mean considering all rats plotted on a session-by-session basis. Data is presented for both identification **(a)** and categorization task **(b)**. Depicted are all the sessions used in the analysis. Solid black lines are the mean and shaded grey the 95% c.i. for fitted bias over all rats and trials within a session. Each session was conducted on a different day.

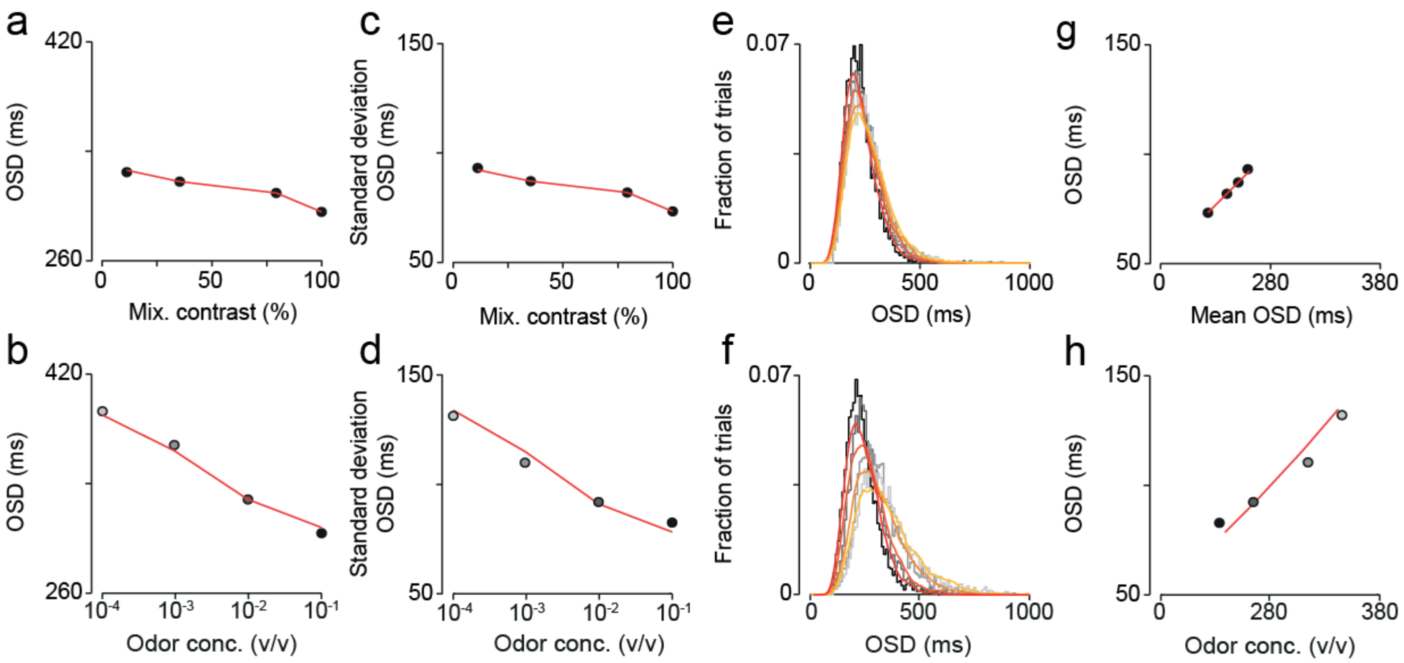


**Figure S21.** Fitting of reaction times to a drift-diffusion model. We fitted the mean and variance of RT to a drift-diffusion model (DDM), where the only parameter allowed to vary across stimuli was the mean drift rate. **(a, b)** Mean RT of a super-rat (all trials from 4 rats pooled together) as a function of mixture contrast (a) or odor concentration (b). Red line corresponds to DDM fit. Dot shading represents odor concentration, with highest concentration corresponding to the darkest shade. **(c, d)** Standard deviation of RT as a function of mixture contrast (c) or odor concentration (d). Red line corresponds to DDM fit. **(e, f)** Histograms of RT (10 ms bin; gray-scale lines) for the different mixture contrast (e) and odor concentration (f) stimuli. Colored lines correspond to the predictions of the DDM (smoothed with an averaging filter, window size = 50 ms). Line shading represents stimulus difficulty, with easier stimuli corresponding to the darkest shade. **(g, h)** Standard deviation of RT as a function of mean RT for the different mixture contrast (g) and odor concentration (h) stimuli. Red line corresponds to DDM fit.

**Supplementary Table 1.** Parameters obtained for best fitting models of the data considering mean psycho- and chrono-metric functions.

| | Parameters | | | | | | | | | | | | | | | |
|---|---|---|---|---|---|---|---|---|---|---|---|---|---|---|---|---|
| **Model** | $\theta$ | $\theta_{slo}$ | $\theta_{dec}$ | $t_{nd}$ | k | $\beta$ | $\lambda$ | $\alpha$ | $\alpha_w$ | $\alpha_z$ | $\sigma_w$ | $\sigma_{wb}$ | $\sigma_d$ | $\sigma_b$ | $p_l$ | $p_b$ |
| DDM | 0.38 | - | - | 0.83 | 24.92 | 0.36 | - | - | - | - | - | - | - | - | - | - |
| DDM + lapses | 0.14 | - | - | 0.31 | 137.35 | 0.41 | - | - | - | - | - | - | - | - | 0.07 | 0.59 |
| DDM + lapses +collapsing bound | 0.15 | -0.12 | - | 0.31 | 151.98 | 0.43 | - | - | - | - | - | - | - | - | 0.09 | 0.59 |
| Bayes-DDM + lapses | 0.09 | - | - | 0.32 | 608.31 | 0.58 | 0.97 | - | - | - | - | - | 0.13 | - | 0.05 | 0.65 |
| Bayes-DDM + lapses + collapsing bound | 0.15 | 0.45 | - | 0.28 | 354.98 | 0.62 | 0.95 | - | - | - | - | - | 0.13 | - | 0.05 | 0.62 |
| Bayes-DDM + lapses + bias | 0.24 | - | - | 0.30 | 343.42 | 0.57 | 0.96 | - | - | - | - | - | 0.13 | 86.00 | 0.05 | 0.68 |
| Full Bayes-DDM | 0.40 | -0.54 | - | 0.28 | 270.47 | 0.57 | 0.96 | - | - | - | - | - | 0.16 | 80.75 | 0.05 | 0.66 |
| Random weights + collapsing bound + bias + lapses | 0.41 | -1.02 | - | 0.28 | 245.91 | 0.57 | 0.91 | - | - | - | 0.17 | - | - | 80.00 | 0.07 | 0.53 |
| Random weights + collapsing bound | 0.37 | 0.19 | - | 0.26 | 270.52 | 0.62 | - | - | - | - | 0.16 | 1.03 | - | - | 0.08 | 0.55 |
| Delta rule + lapses | 0.12 | - | - | 0.31 | 135.95 | 0.38 | - | 0.10 | - | - | - | - | - | - | 0.05 | 0.66 |
| Delta rule + collapsing bounds | 0.07 | 1.00 | - | 0.32 | 490.18 | 0.52 | - | 0.11 | - | - | - | - | - | - | 0.14 | 0.59 |
| Delta rule + bias + lapses | 0.37 | - | - | 0.27 | 252.57 | 0.61 | - | 0.03 | - | - | - | - | - | - | 0.06 | 0.44 |
| Delta rule + collapsing bounds + bias + lapses | 1.19 | -4.02 | - | 0.23 | 257.06 | 0.64 | - | 0.08 | - | - | - | - | - | - | 0.05 | 0.50 |
| RL-DDM | 0.50 | - | 7.81 | 0.29 | 626.00 | 0.71 | - | - | 0.04 | 0.03 | - | - | - | - | 0.07 | 0.57 |

Presented results were obtained while simultaneously fitting both categorization and identification task. The presented models where fit to maximize the log-likelihood of the mean curves of all rats combined (See Experimental Procedures on model fitting for more details). This analysis was also done for all rats and interleaved condition (not shown).

**Supplementary Table 2.** Parameters obtained for best fitting models of the data considering mean and sequential (dependent on previous trial outcome) psycho- and chrono-metric functions.

| | Parameters | | | | | | | | | | | | | | | |
|---|---|---|---|---|---|---|---|---|---|---|---|---|---|---|---|---|
| **Model** | **$\theta$** | **$\theta_{slo}$** | **$\theta_{dec}$** | **$t_{nd}$** | **k** | **$\beta$** | **$\Lambda$** | **$\alpha$** | **$\alpha_w$** | **$\alpha_z$** | **$\sigma_w$** | **$\sigma_{wb}$** | **$\sigma_d$** | **$\sigma_b$** | **$p_l$** | **$p_b$** |
| DDM | 0.91 | - | - | 1.00 | 9.38 | 0.35 | - | - | - | - | - | - | - | - | - | - |
| DDM + lapses | 0.16 | - | - | 0.31 | 152.06 | 0.43 | - | - | - | - | - | - | - | - | 0.10 | 0.57 |
| DDM + lapses +collapsing bound | 0.15 | 0.12 | - | 0.31 | 152.06 | 0.43 | - | - | - | - | - | - | - | - | 0.09 | 0.60 |
| Bayes-DDM + lapses | 0.12 | - | - | 0.31 | 625.97 | 0.60 | 0.97 | - | - | - | - | - | 0.13 | - | 0.05 | 0.64 |
| Bayes-DDM + lapses + collapsing bound | 0.15 | 0.47 | - | 0.27 | 355.00 | 0.62 | 0.95 | - | - | - | - | - | 0.10 | - | 0.09 | 0.58 |
| Bayes-DDM + lapses + bias | 0.24 | - | - | 0.30 | 357.00 | 0.59 | 0.97 | - | - | - | - | - | 0.11 | 86.00 | 0.05 | 0.62 |
| Full Bayes-DDM | 0.41 | -0.59 | - | 0.29 | 268.46 | 0.58 | 0.97 | - | - | - | - | - | 0.16 | 80.60 | 0.04 | 0.69 |
| Random weights + collapsing bound + bias + lapses | 0.41 | -1.02 | - | 0.28 | 245.91 | 0.57 | 0.91 | - | - | - | 0.17 | - | - | 80.00 | 0.07 | 0.55 |
| Random weights + collapsing bound | 0.37 | 0.19 | - | 0.26 | 270.49 | 0.62 | - | - | - | - | 0.16 | 1.05 | - | - | 0.09 | 0.59 |
| Delta rule + lapses | 0.12 | - | - | 0.31 | 134.25 | 0.38 | - | 0.11 | - | - | - | - | - | - | 0.05 | 0.54 |
| Delta rule + collapsing bounds | 0.10 | 1.00 | - | 0.32 | 496.01 | 0.58 | - | 0.13 | - | - | - | - | - | - | 0.10 | 0.60 |
| Delta rule + bias + lapses | 0.38 | - | - | 0.28 | 580.27 | 0.72 | - | 0.03 | - | - | - | - | - | - | 0.08 | 0.45 |
| Delta rule + collapsing bounds + bias + lapses | 1.27 | -4.96 | - | 0.23 | 259.00 | 0.64 | - | 0.08 | - | - | - | - | - | - | 0.05 | 0.50 |
| RL-DDM | 0.51 | - | 7.75 | 0.29 | 626.00 | 0.71 | - | - | 0.04 | 0.13 | - | - | - | - | 0.05 | 0.49 |

Presented results where obtained while simultaneously fitting both categorization and identification task. The presented models where fit to maximize the log-likelihood of the mean and conditional curves of all combined rats (See Experimental Procedures on model fitting for more details). This analysis was also done for all rats (not shown).
